# Larger Numbers Can Impede Adaptation in Asexual Populations despite Entailing Greater Genetic Variation

**DOI:** 10.1101/076760

**Authors:** Yashraj D. Chavhan, Sayyad Irfan Ali, Sutirth Dey

**Affiliations:** Indian Institute of Science Education and Research (IISER) Pune, Dr. Homi Bhabha Road, Pashan, Pune, Maharashtra, 411008, India.; Indian Institute of Science, Bangalore, Karnataka, 560012, India.

**Keywords:** Predicting adaptation, effective population size, experimental evolution, extent of adaptation, population bottlenecks, adaptive size, adaptive dynamics

## Abstract

Periodic bottlenecks play a major role in shaping the adaptive dynamics of natural and laboratory populations of asexual microbes. Here we study how they affect the ‘Extent of Adaptation’ (*EoA*), in such populations. *EoA*, the average fitness gain relative to the ancestor, is the quantity of interest in a large number of microbial experimental-evolution studies which assume that for any given bottleneck size (*N*_0_) and number of generations between bottlenecks (*g*), the harmonic mean size *(HM=N_0_g*) will predict the ensuing evolutionary dynamics. However, there are no theoretical or empirical validations for *HM* being a good predictor of *EoA*. Using experimental-evolution with *Escherichia coli* and individual-based simulations, we show that *HM* fails to predict *EoA* (i.e., higher *N_0_g* does not lead to higher *EoA*). This is because although higher *g* allows populations to arrive at superior benefits by entailing increased variation, it also reduces the efficacy of selection, which lowers *EoA*. We show that *EoA* can be maximized in evolution experiments by either maximizing *N_0_* and/or minimizing *g*. We also conjecture that *N_0_/g* is a better predictor of *EoA* than *N_0_g*. Our results call for a re-evaluation of the role of population size in predicting fitness trajectories. They also aid in predicting adaptation in asexual populations, which has important evolutionary, epidemiological and economic implications.

## Introduction

Population size is a key demographic parameter that affects several ecological and evolutionary processes including the rate of adaptation (Desai et al. 2007; Desai and Fisher 2007; Gerrish and Lenski 1998; Lanfear et al. 2014; Samani and Bell 2010; Wilke 2004), efficacy of selection (Petit and Barbadilla 2009), organismal complexity (LaBar and Adami 2016), fitness decline (Katju et al. 2015), repeatability of evolution (Lachapelle et al. 2015; Szendro et al. 2013; Vogwill et al. 2016), etc. Interestingly though, what constitutes a useful measure of population size for predicting evolutionary outcomes often depends on the ecological/evolutionary question being addressed and the population-genetics quantity in question (Charlesworth 2009). Consequently, it is crucial to use the relevant measure of population size while constructing or empirically validating any evolutionary theory.

Experimental evolution using asexual microbes has been one of the key tools in validating several tenets of evolutionary theory ((Kassen 2014), reviewed in (Kawecki et al. 2012)). Most such studies deal with populations that face regular and periodic bottlenecks during their propagation (Kawecki et al. 2012). The absolute population size keeps changing regularly because of these periodic bottlenecks. Therefore, in order to make predictions and claims based on population size in such experiments, it is important to define a proper measure of population size depending upon the question of interest (Charlesworth 2009; Kawecki et al. 2012; Lanfear et al. 2014; Wang et al. 2016).

Previous theoretical studies have shown that the harmonic mean of population size *(HM)* over time acts as the measure of population size that can explain and predict the fixation probabilities of beneficial mutations in such systems (Patwa and Wahl 2008; Wahl and Gerrish 2001). Specifically, if a population grows from size *N_0_* to *N_f_* via binary fissions within a growth phase, and is diluted back periodically to *N_0_* by random sampling at the end of the growth phase, then the relevant population-size measure for fixation probabilities is given by *N_e_* ≈ *N_0_*\**log_2_(N_f_/N_0_)* = *N_0_g*, where *g* refers to the number of generations between successive bottlenecks and *N_0_g* is the harmonic mean size (Lenski et al. 1991). From an evolutionary perspective, periodic bottlenecks play two opposite roles in such experiments. On the one hand, harsher bottlenecks (entailed by higher *g*) reduce the probability that a given beneficial mutation would fix due to sampling errors during the bottleneck. On the other hand, higher values of *g* also imply an increase in *N_f_*, which causes an increase in mutational opportunities (binary fissions) during exponential growth (Wahl et al. 2002). This is expected to increase the total supply of mutations that would survive drift, which in turn should increase the raw material available for evolution. It has been predicted that exponential growth between *N_0_* and *N_f_* influences fixation probabilities more than the elimination by sampling (Heffernan and Wahl 2002). More nuanced and complex measures of population size (Campos and Wahl 2009, 2010) also suggest that adaptation rates in terms of fixation probabilities would have a positive relationship with *N_0_* and *g*, the two population size parameters amenable to experimental manipulation.

Unfortunately, most experimental evolution studies with serially bottlenecked asexual populations do not focus on the fixation probabilities of beneficial mutants. Instead, they are interested in the average amount of fitness gained with respect to the ancestor at a given time (we call this quantity the extent of adaptation, *EoA*) during the course of evolution (De Visser and Rozen 2005; Desai et al. 2007; Lachapelle et al. 2015; Lenski et al. 1991; Rozen et al. 2008; Samani and Bell 2010). Several experimental studies, dealing with quantities akin to *EoA* for periodically bottlenecked asexual populations, have used *HM (*=*N_0_g)* for quantifying the evolutionarily relevant (i.e. predictive of the magnitude of evolutionary response) population size (De Visser and Rozen 2005; Desai et al. 2007; Lachapelle et al. 2015; Lenski et al. 1991; Rozen et al. 2008; Samani and Bell 2010). However, to the best of our knowledge, there is no theoretical basis or empirical justifications (Y. Raynes et al. 2014) for this usage of *HM*. Here we use a combination of agent-based simulations and long-term evolutionary experiments using *Escherichia coli* to investigate the interplay of *N_0_* and *g* in shaping the *EoA* of asexual populations. Since *HM* has been widely used by experimentalists in the context of *EoA*-like quantities, we begin by testing the suitability of *HM* as a predictor of *EoA*. We show that populations with similar values of *N_0_g* can have markedly different *EoA* trajectories, and this result applies to both real (bacterial) as well as simulated populations. Secondly, we demonstrate that although increasing the value of *g* (making the periodic bottleneck harsher) promotes adaptation through an increased supply of variation, it also reduces the efficacy of selection which impedes adaptation by restricting the spread of large-effect beneficial mutations. When these two opposing aspects of bottlenecks are considered together, counterintuitively, *EoA* turns out to have a negative relationship with *g*. Thirdly, we show that populations with similar *HM* can not only have different fitness trajectories but can also differ markedly in terms of how frequency-distribution of fitness amongst individuals changes during adaptation. Finally, we show that, for a given mutation rate, *N_0_/g* can be a better predictor of *EoA* trajectories, *i.e*., populations with similar *N_0_/g* have similar fitness trajectories and populations with higher *N_0_/g* adapt faster. Our findings thus introduce a new way of thinking about the relationship between population size and adaptive trajectories.

Our approach differs from previous studies in two important ways. First, unlike many studies (Campos and Wahl 2009, 2010; Heffernan and Wahl 2002; Wahl and Gerrish 2001) we focus on how *EoA* (and not long-term fixation probabilities) is shaped by bottleneck size (*N_0_) and bottleneck ratios (*N_0_/N_f_*). This makes our study directly relevant to a rich body of microbial experimental evolution literature ((De Visser and Rozen 2005; Desai et al. 2007; Lachapelle et al. 2015; Lenski et al. 1991; Rozen et al. 2008; Samani and Bell 2010), reviewed in (Kawecki et al. 2012)). Second, many previous theoretical studies on periodically bottlenecked systems (where *N_f_* = *N*_0_2^g^*), assume that the culture volume (and therefore *N_f_*) is a constant, and then go on to explore what value of *N_0_* or *g* leads to the minimum loss of variation during bottlenecks and/or in the long run (Campos and Wahl 2009, 2010; Heffernan and Wahl 2002; Wahl et al. 2002; Wahl and Gerrish 2001; Wahl and Zhu 2015). In our simulations, we remove this restriction and seek to compare loss of variation in those cases where both *N_0_* and *N_f_* can be different (e.g. between a population grown in 50 ml of medium versus one grown in (say) 1 ml of medium). Clearly, it is possible to have two populations with very different *N_0_* and *N_f_* values that can nevertheless have similar values of *N_0_g*. One of the questions that we investigate is whether such populations have similar fitness trajectories or not. Thus, our results make it possible to compare the expected *EoA* across experimental studies that employ similar environments but different culture volumes, which is a rather common scenario in experimental evolution studies (Lachapelle et al. 2015; Y. Raynes et al. 2014; Yevgeniy Raynes et al. 2012; Rozen et al. 2008; Samani and Bell 2010).

## Methods

### Experimental evolution, assays and statistical analysis

Here we present a brief description of the experimental protocol, relegating the details to Supplementary Methods. Our primary aim was to investigate if a commonly used measures of population size in experimental evolution, namely harmonic mean (*N_0_g*), could predict *EoA* trajectories. We also wanted to see if populations with similar values of *N_f_* have similar *EoA*. To this end, we experimentally evolved three different population regimens (LL, SL, and SS) in Nutrient broth containing a sub-lethal cocktail of three antibiotics (Norfloxacin, Rifampicin and Streptomycin) for ~380 generations in batch culture. Each regimen consisted of 8 independently evolving replicate populations, all of which were started from a single *Escherichia coli* MG 1655 colony. The three population regimens were propagated at different bottleneck sizes: LL faced lenient bottlenecks (1/10), whereas SS (1/10^4^) and SL (1/10^6^) experienced much harsher bottlenecks. LL and SL were grown at larger culture volumes (100 ml, culture in flasks) than SS (1.5 ml, culture in 24 well-plates). Thus, in terms of *N_f_*, LL = SL ≫ SS but in terms of *N_0_g* LL ≫ SL = SS (see Table S1 for the values of these parameters). The *EoA*-trajectories of the three population types were reconstructed by assaying the maximum population-wide growth rates (*R*) and carrying capacities (*K*) of each replicate at different time-points during evolution following standard protocols (Karve et al. 2015, 2016), the details of which can be found in Supplementary Methods. *K* of a population was defined as the maximum OD value attained over a period of twenty four hours (the highest value in the sigmoidal growth curve) (Karve et al. 2016; Novak et al. 2006). *R* was estimated as the maximum slope of the growth curve over a running window of four OD readings (each window spanning one hour) (Karve et al. 2015, 2016; Vogwill et al. 2016).

To analyze the data, we performed separate repeated measures (RM) ANOVA for *K* and *R*. “Regimen-type” (LL/SL/SS) was treated as the categorical factor, and TIME (9 time-points) as the repeated measures factor. We also included an interaction of Regimen-type and TIME in our model to determine if the fitness trajectories of the regimes were significantly different from each other. Furthermore, to compare the fitness values at each time point, we used a nested-design ANOVA with “regimen-type” (SS, SL or LL, fixed factor) and “replicate-line” (1–8, random factor, nested in regimen-type). We used Holm-Šidàk correction (Abdi 2010) for controlling the family-wise error rates. For all ANOVAs where there was a significant effect of “Regimen-type” after the Holm-Šidàk correction, we used Tukey’s HSD to compare pairwise differences between LL, SL, and SS (See Supplementary Methods for details and rationale).

### Simulations of microbial evolution

Any difference between the three regimens in our experiment can, in principle, be due to some idiosyncratic properties of the experimental organism (*E. coli*) or potential differences between the selection environments in flasks and plates. In order to account for that possibility and enhance the generalizability of our results, we used an Individual Based Model (IBM) to simulate bacterial growth under resource-limited conditions (Wahl et al. 2002). Except for differences in the amount of resources, our model contained no other parameters specific to *E. coli* or related to differences in culture conditions. Thus, in terms of differences between the *EoA* of the regimens, if the model output matched the empirical observations then our results were likely to be applicable for other asexual systems. Treating our experiment as a case-study, we used our model to investigate if our results were generalizable.

Our simulations start with a nearly clonal distribution of fitness effects. In our model, an individual bacterium was characterized by three principal parameters: efficiency, threshold, and body-mass. The simulation (coded in the C programming language) began with a fixed amount of resources available in the environment, utilized by the bacteria for growth. A typical individual was represented by an array that specified three principal parameters: (1) Bodymass, (2) Efficiency, and (3) Threshold. Efficiency and Threshold were the only two evolvable parameters. Bacteria consumed resources in an iterative and density-dependent manner. The parameter *Bodymass_i_* of the i^th^ individual represented how big the particular individual was during a given iteration. Its efficiency *(K_eff_i_)* specified how much food it assimilated per iteration. If *population size/ K_eff_i_ < 1, 10(1 - (population size/ K_eff_i_))* units were added to *Bodymass_i_*. Otherwise, *Bodymass_i_* remained unchanged. *Bodymass_i_* increased with cumulative assimilation. When *Bodymass_i_* becomes greater than or equal to *thres_i_* (its threshold parameter), the individual *i* underwent binary fission and divided into two equally sized daughter individuals. Each fission event had a fixed probability of giving rise to mutations based on a mutation rate that remained constant for all individuals in the population. *K_eff_i_* and *thres_i_* mutate independently, and were the only two parameters that could undergo mutation. The mutated value was drawn from a static normal distribution with the frequency of deleterious mutations being much higher than that of beneficial mutations, which is in line with experimental observations (See Table S3; Kassen and Bataillon, 2006; Eyre-Walker and Keightley, 2007). The distribution of mutational effects remained fixed throughout the simulation (Kassen and Bataillon, 2006) due to which, *EoA* was expected to eventually approach a plateau. When the population ran out of resources (once the amount of body-mass accumulated per unit time by the population went below a pre-decided threshold so that the sigmoidal curve reached a plateau), it was sampled according to the sampling ratio being studied. The above process was repeated for 400 generations, where each generation represented two-fold growth in population size (see Supplementary Methods for a more detailed description of the model). We also checked if our model met several intuitive theoretical predictions that had not been coded directly (see Supplementary data (SD1); also see Figs. S1, S2, and S3).

Density-dependent growth, clonal interference (CI), the presence of deleterious mutations, the presence of variable fitness effects of mutations, etc. are some key features that are instrumental in shaping the adaptive dynamics of periodically bottlenecked asexual populations (Patwa and Wahl 2008; Sniegowski and Gerrish 2010). Unfortunately, the complex interactions of so many features are difficult to capture in analytical models (Sniegowski and Gerrish 2010). Consequently, previous theoretical studies have been forced to make simplifying assumptions like the absence of deleterious mutations (Desai and Fisher 2007; Wahl and Gerrish 2001), constancy of beneficial mutational effects (Desai and Fisher 2007), constancy of *N_f_* (Campos and Wahl 2009, 2010; Wahl and Gerrish 2001; Wahl and Zhu 2015), the presence of discrete generations (Campos and Wahl 2009, 2010; Desai and Fisher 2007), etc. (see Table S2 for details). Our IBM avoids these simplifying assumptions, which might explain why some of the features captured by our model have not been reported earlier. Moreover, our study is in the context of *EoA*, while most of the earlier studies have investigated fixation probabilities.

## Results

### *HM* failed to predict and explain the *EoA* trajectories of experimental populations

RM ANOVA on all three regimens indicated a significant Regimen-type × TIME interaction for both *K* (F_16, 168_ = 5.72; p < 0.000001) and *R* (F_16, 168_ = 7.306; p < 0.000001). However, in principle, this interaction could be driven by the fact that the LL populations had a much larger increase in *K* and *R* compared to the SL and SS populations. Since our primary interest was to check whether the SL and SS populations differed in terms of these two fitness measures, we repeated the RM ANOVA for only these two regimens and again found a significant Regimen-type × TIME interaction for both *K* (*F*_8, 112_ = 2.070; *p* = 0.0446) and *R* (*F*_8, 112_ = 3.594; *p* = 0.000948). Since the interaction term was significant, we chose not to interpret the main effects of Regimen-Type or TIME.

Individual ANOVAs showed that the *EoA* of SS was greater than that of SL at 5/6 and 4/5 time-points which had significant difference in terms of *K* (Fig. 1a) and *R* (Fig. 1b). The *p*-values and the *F*-values (with corresponding *df*) for each time-point for *K* and *R* are presented in Tables S4 and S5 respectively. Thus, particularly during the last two thirds of the evolution experiment, the *EoA* of SS was consistently higher than that of SL. The effect sizes (Cohen’s *d* (Cohen 1988)) of *EoA* differences between SL and SS were found to be either medium or large (with the majority being large effects; see Table S6) for several points on the *EoA* trajectory. Thus, similar *HM* can give rise to fairly different adaptive trajectories. This observation is consistent with recent empirical findings that question the validity of harmonic mean as an evolutionarily relevant population size (Y. Raynes et al. 2014). Surprisingly, SS had a larger overall *EoA* than SL despite having lower *N_f_*. Interestingly, despite having similar *N_f_*, LL typically had much larger *EoA* than SL, which is explainable by the fact that the latter regimen suffered more severe bottlenecks. This shows that similar *N_f_* does not lead to similar *EoA* if the bottleneck ratios are different.

**Fig. 1.**
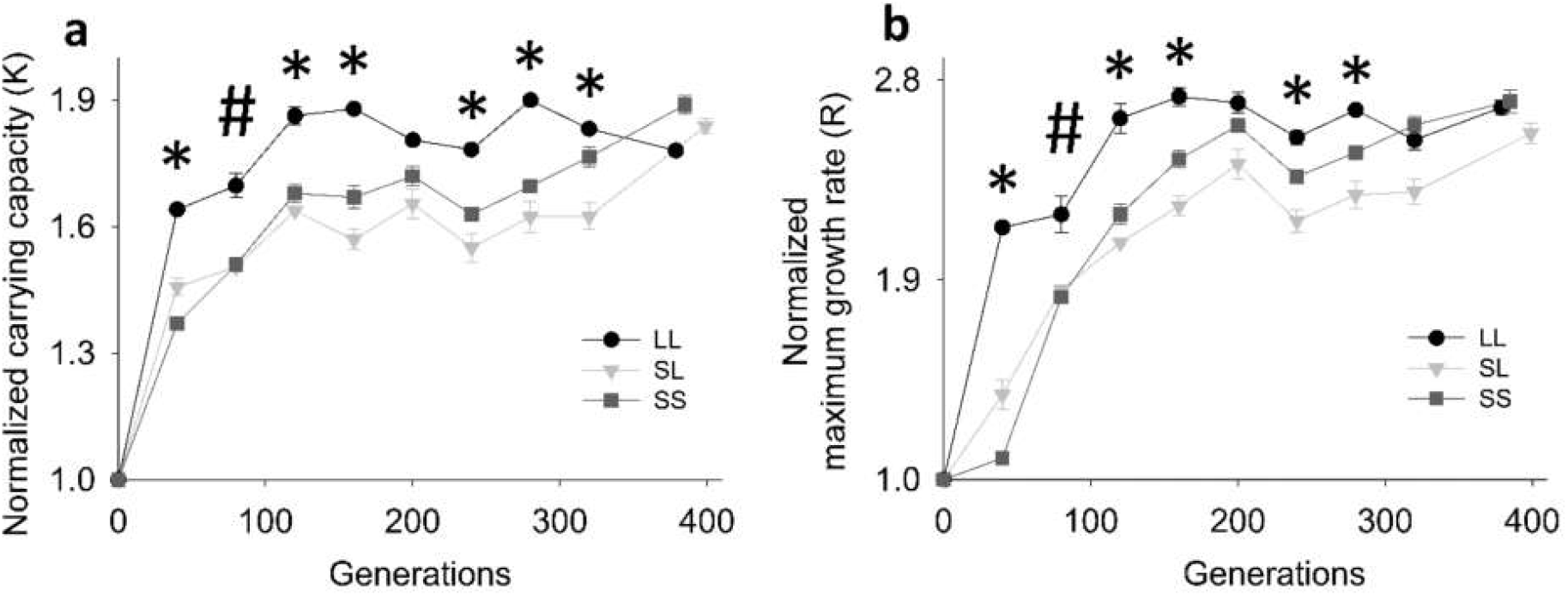
Experimental *EoA* trajectories in terms of carrying capacity and maximum growth rate. (**a**) *EoA* of carrying capacity *(K)*. (**b**) *EoA* of maximum growth rate *(R)*. Data points show mean ± SEM for 8 replicates. * refers to cases when all three pairwise differences (LL-SL, LL-SS, and SL-SS) are significant (Tukey post hoc *p* < 0.05). # refers to significant difference across LL-SL and LL-SS, but not SL-SS (See Tables S4 and S5). SS and SL have markedly different adaptive trajectories despite having similar harmonic mean population sizes.

In summary, *HM* failed to predict the *EoA* trajectories of our experimental populations as, in spite of having similar values of *N_0_g*, the SL and SS regimens had markedly different adaptive *(EoA)* trajectories for *K* (Fig. 1a) as well as *R* (Fig. 1b).

### Simulations also revealed that *HM* fails to predict adaptive trajectories

We simulated evolution in populations with similar *HM* but with different values of *N_0_* and *g*, such that the product (*N_0_g*) remained constant. If *HM* (=*N_0_g*) were a good predictor of how much a population is expected to adapt, then these three treatments were expected to show similar *EoA*. This was not found to be the case for both *K* (Fig. 2a) and *R* (Fig. 2b), which was consistent with our experimental observations of *EoA* trends in SL and SS (Fig. 1; also see Supplementary Methods and Fig. S4). XX’, SS’, and SL’ were also found to be remarkably different in terms of the adaptive increase in average efficiency of individuals (Fig. S4a). Interestingly, populations with similar *HM* were also found to differ in terms of the frequency distributions of the efficiency parameters amongst their constituent individuals (Fig. S5). To elucidate why *N_0_g* could not explain *EoA* trajectories, we determined how *EoA* varied with *N_0_* and *g*, independently.

**Fig. 2.**
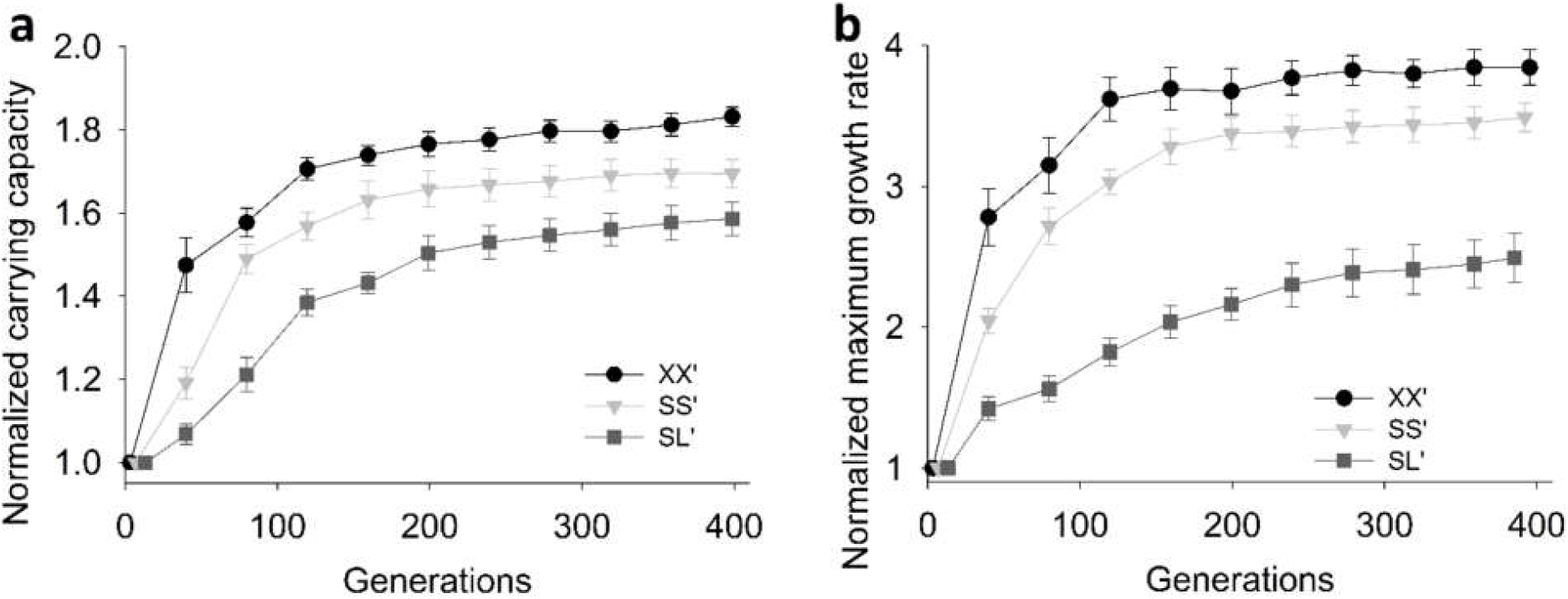
Simulations: Adaption in three populations with similar harmonic mean size. Data points show mean *EoA* ± SEM for 8 replicates. (**a**) Adaptation in terms of normalized carrying capacity *(K)*. (**b**) Adaptation in terms of normalized maximum growth rate *(R)*. XX’, SS’ and SL’ had similar harmonic mean sizes and represent lenient, medium and harsh bottlenecks with *N_0_* ≈ 3.6×10^3^, 1.8×10^3^, 9×10^2^ and bottleneck ratio of 1/10; 1/10^2^,1/10^4^ respectively. These simulations suggest that populations with similar harmonic mean size can have markedly different *EoA* trajectories.

### *EoA* varied positively with *N_0_* but negatively with *g*

If *N_0_g* were a good measure of the population size that has a positive relationship with *EoA*, then increasing either *N_0_* or *g* or both should lead to greater *EoA*. We tested this intuitive prediction via simulations using several combinations of *N_0_* and *g*, spanning four orders of magnitude for both *N_0_* and the sampling ratio (*N_0_/N_f_*). Although *EoA* was found to increase with greater *N_0_* (Fig. 3a; also see Fig. S6), the relationship between *EoA* and *g* turned out to be negative (Fig. 3b; also see Fig. S7) which was reflected in terms of both individual-level (Fig. 3b—in terms of efficiency) and population-level (Fig. S8—in terms of *R*) fitness parameters. The latter result implied that larger values of *N_f_* impeded adaptation in populations when the population size during the bottleneck *(N_0_)* was held constant. The nature (sign) of this relationship between *EoA* and *g* was found to be robust to changes in mutation rate over a 100-fold range in our simulations (Fig. S10). A negative relationship between *EoA* and *g* is particularly surprising because, in populations with similar *N_0_*, increase in *g* is expected to lead to an increase in the available variation. All else being equal, this should have led to greater *EoA*. Since that was not the case, we went on to check if these slowly adapting populations (with similar *N_0_* but higher *g*) were limited, qualitatively and/or quantitatively, by the availability of variation.

**Fig. 3.**
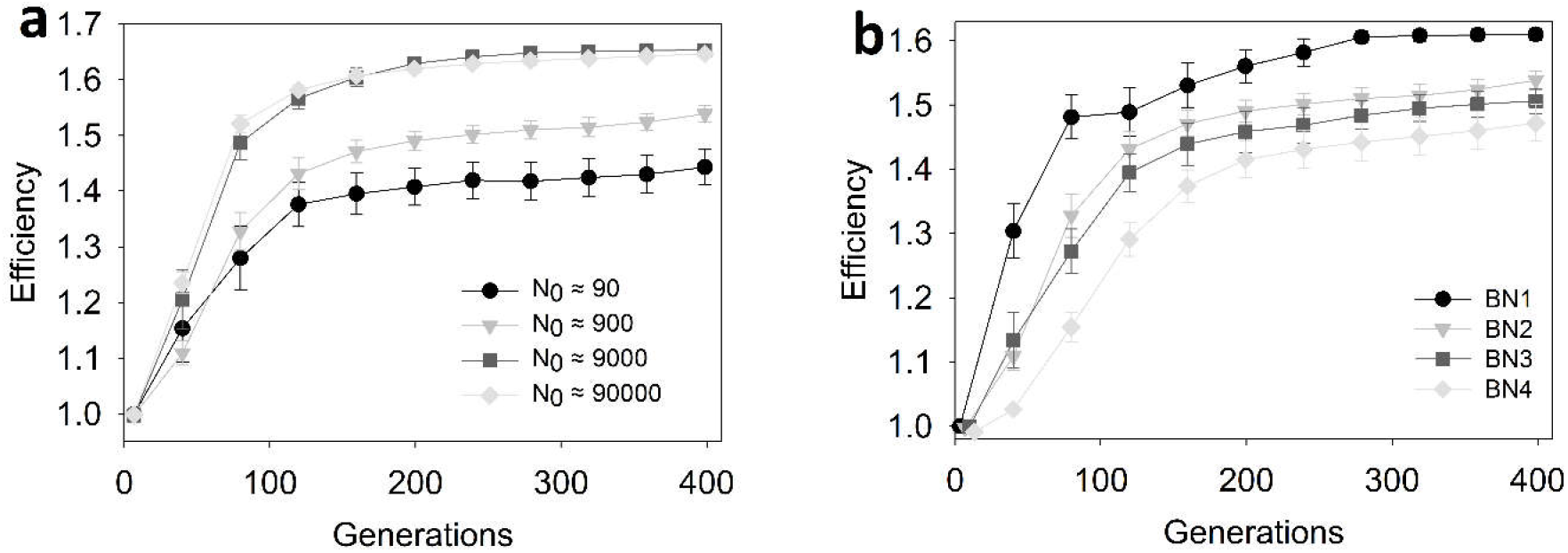
Simulations: The relationship of *EoA* (expressed in terms of efficiency) with *N_0_* and *g*. Data points show mean ± SEM; 8 replicates. The populations shown in (**a**) had the same bottleneck ratio (1/10^2^) but different bottleneck sizes (= *N_0_*). *EoA* varies positively with *N_0_*. On the other hand, the populations shown in (**b**) had identical bottleneck size (= *N_0_)* but different bottleneck ratios (reflected by different values of *g*. Bottleneck ratios: BN1: 1/10 (*g* = 3.32); BN2: 1/10^2^ (*g* = 6.64); BN3: 1/10^3^ (*g* = 9.96); BN4:1/10^4^ (*g* = 13.28). *EoA* varies negatively with *g*. Also see Figs. S6, S7 and S8.

### The quantitative availability of beneficial traits could not explain why *EoA* varied negatively with *g*

Consider SM1 and SM4, treatment regimens which had similar starting population size (*N_0_*) after the first bottleneck but had *g* values of 3.32 and 13.28 respectively (SM refers to sampling ratio, expressed in terms of log(10) (see Figs. 4 and 5). SM1 grew to a final size of 10N0 in one growth phase (*i.e*., before bottleneck), while SM4 grew to 10^4^*N_0_*. Consequently, SM1 faced a periodic bottleneck of 1/10 whereas SM4 was sampled 1/10^4^. Since SM4 experienced approximately 278 times more fission events than SM1 per evolutionary generation, the former was expected to undergo more mutations and consequently show more variation. Moreover, SM4 was also expected to arrive at very large-effect benefits that were so rare that the probability of SM1 stumbling upon them was vanishingly low due to its lower mutational supply. As expected, compared to SM1, SM4 had a greater within-population coefficient of variation in terms of efficiency values (Fig. 4) and therefore was not limited by the supply of variation. To better understand the contributions of phenotypes of different magnitudes to *EoA*, we classified the phenotypes into 50 discrete static classes. We found that SM4 also had a continuous access to highly fit genotypes (Fig. S11a) that were inaccessible to SM1 throughout the simulations. On the basis of these observations, *EoA* can be expected to vary positively with *g* and thus SM4 was expected to be fitter than SM1 at a given point of time in general. However, counterintuitively, SM4 had a consistently lower *EoA* than SM1 (Fig. 4). Evidently, harsher periodic sampling impeded adaptation despite resulting in increased substrate for selection. We also found that although higher *N_f_* allowed SM4 to arrive at extremely rare mutations with very large benefits, these mutations failed to survive the harsh periodic bottlenecks by rising to large enough frequencies (Fig. S12a). In other words, SM4 typically wasted the best mutation explored by it but SM1 almost always conserved it. This explains why arriving at these rare mutations with very large benefits did not make SM4 adapt more than SM1 in a sustained manner. However, this does not explain why the *EoA* in SM4 was consistently lower than that of SM1.

**Fig. 4.**
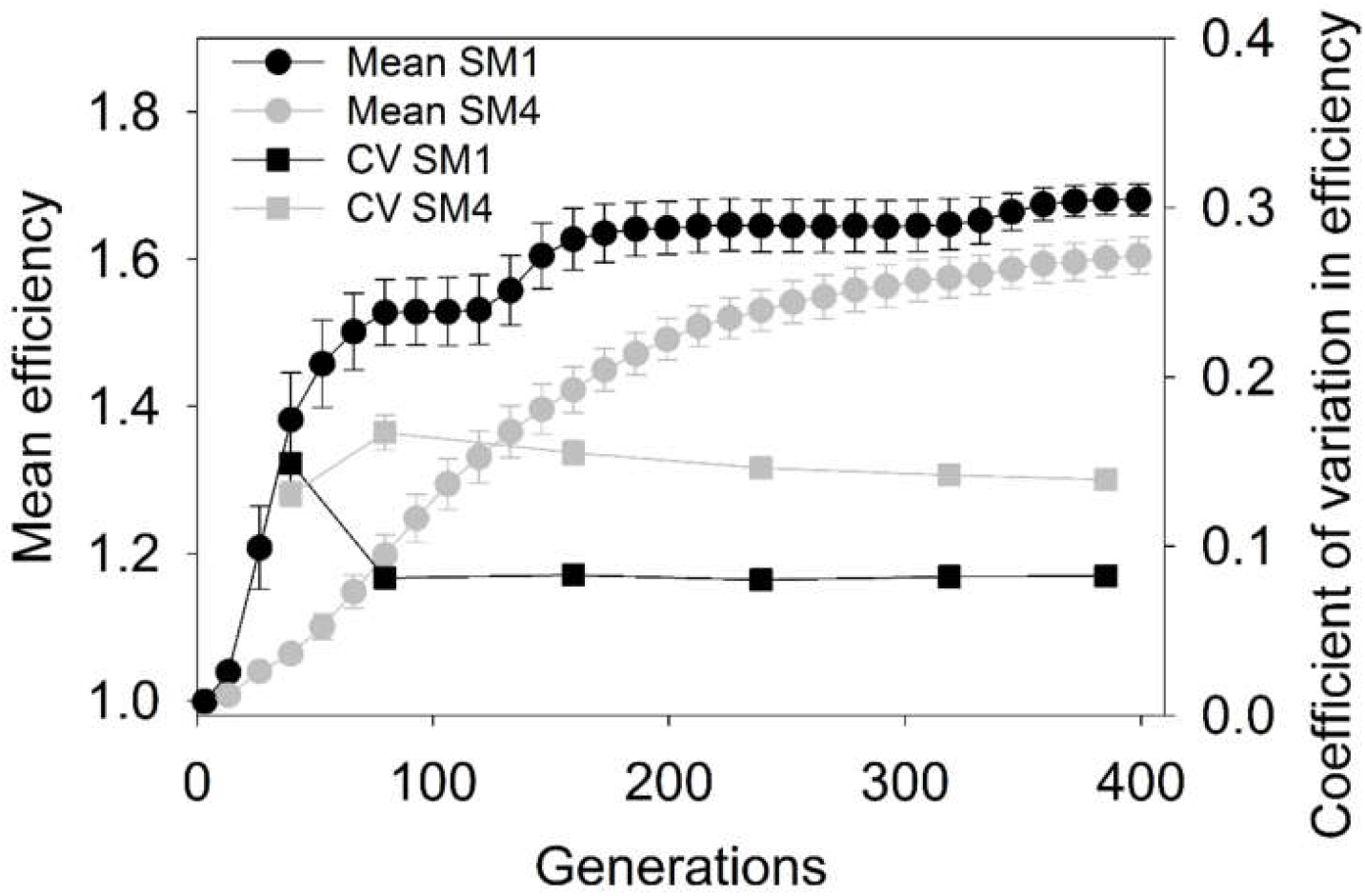
Simulations: Trajectories of efficiency in terms of across-population mean and within-population coefficient of variation. The within-populations coefficient of variation (CV) was computed for each replicate population across its constituent individuals using discrete frequency distributions. The error bars represent SEM (8 replicates). Both SM1 and SM4 had similar bottleneck size (*N_0_* ≈ 900). SM1 experienced a periodic bottleneck of 1/10 whereas SM4 experienced a periodic bottleneck of 1/10^4^. SM4 had a consistently lower *EoA* than SM1 despite having consistently more variation.

### The negative relationship between *EoA* and *g* can be explained in terms of the efficacy of selection

The efficacy of selection in eliminating deleterious mutations and spreading beneficial ones is an important factor that influences the increase of *EoA*. We quantified the inefficacy of selection in increasing *EoA* using the Genetic Load (*GL*), which was defined as: *GL* = *(Best Efficiency – Average Efficiency)/Best Efficiency* (Crow 1958; Rice 2004). The term “Best Efficiency” refers to the highest efficiency value that succeeds in surviving the bottleneck. As discussed earlier, the magnitudes of the highest efficiency values explored by SM4 populations are much greater than those explored by SM1 (Fig. S11a). However, these high-fitness phenotypes of SM4 typically have such low frequencies that they almost always fail to survive bottlenecks and thus do not contribute significantly to the *EoA* (Fig. S12a). Therefore, we defined GL only in terms of the phenotypes that survived the bottlenecks. We found that the Best Efficiency (after bottlenecks) for SM1 was very similar to that of SM4 (Fig. S12b). We note here that the phenotypes that are fitter than the wild type but less fit than the best phenotype also contribute to the GL. Thus, consistently higher GL entails lower contribution of the best phenotype to the *EoA*. Furthermore, if these best phenotypes (with respect to which GL is defined) are similar across populations being compared, consistently lower contribution of the best phenotype to *EoA* would in turn entail slower rise of *EoA*.

SM4 consistently experienced a heavier genetic load than SM1, particularly during the initial phases of evolution (Fig. 5a). This genetic load was constituted largely by phenotypes that are fitter than the wild type ancestor but less fit than the best phenotype (Fig. S13). We labelled the top five occupied fitness classes as the “nose” (*sensu* Desai and Fisher (2007)) and all the classes inferior to the nose as the “lagging chunk.” During the early phases of evolution, the relative contribution of the lagging chunk to the *EoA* was much higher in SM4 than in SM1 (Fig. 5b). In other words, the nose accounted for most of the *EoA* in SM1 but not in SM4 (shown schematically in Figs. 5c and 5d). Thus, compared to SM1, the best phenotype of a typical SM4 population needed to outcompete many more phenotypes (present in sizable frequencies) that were superior to the wild-type but inferior to itself. This suggests that the efficacy of selection was higher in SM1 than in SM4, which in turn explains the faster increase in *EoA* in the former. As selection proceeded, the GL of SM4 reduced greatly by generation 360 (Fig. 5a). This resulted in similar contributions of the respective noses to the overall *EoA* in SM1 and SM4 (Figs. 5e and 5f).

**Fig. 5.**
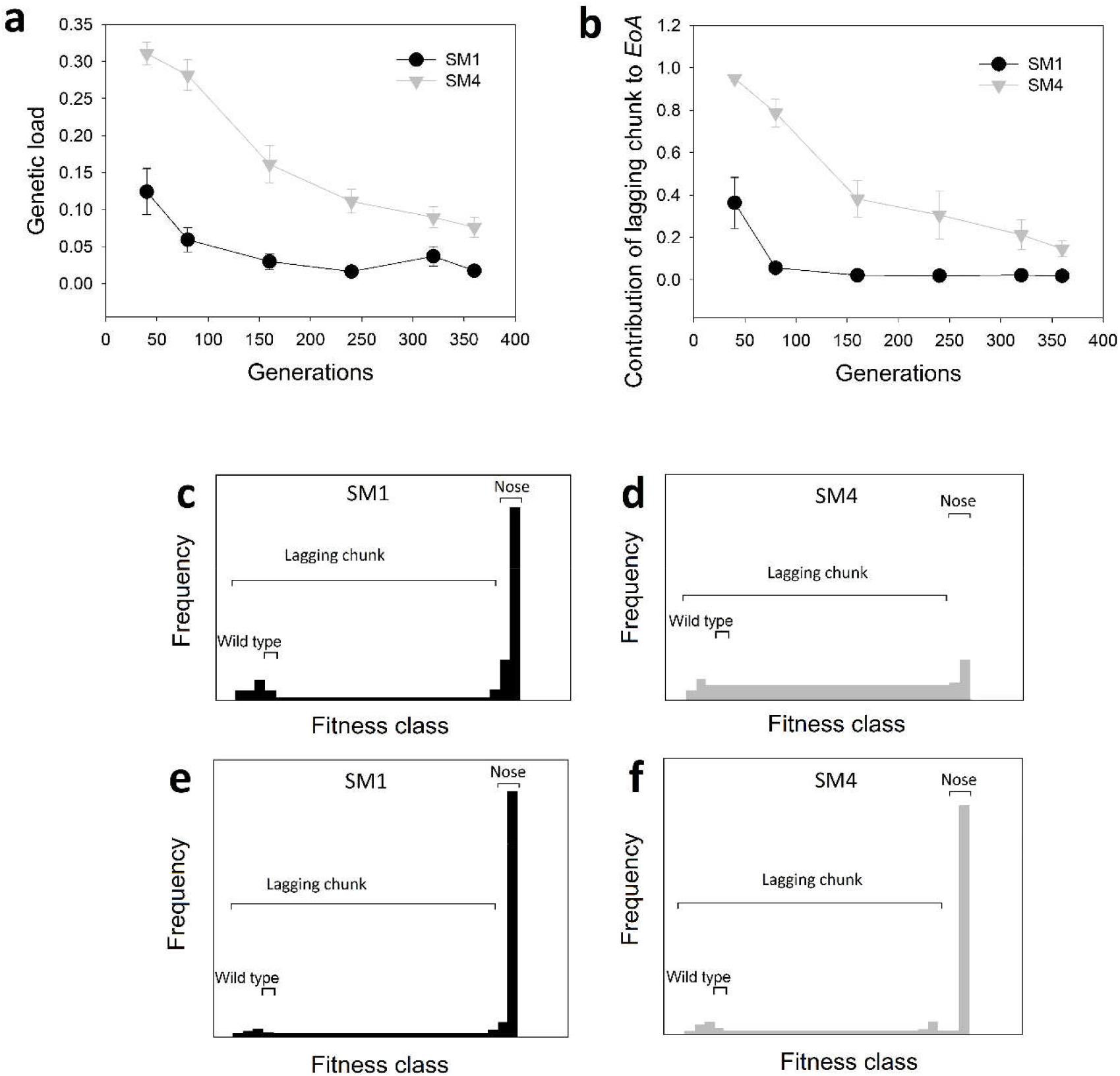
The efficacy of selection in SM1 was more than that in SM4. (**a**) SM1 consistently experienced a much lower genetic load than SM1 (the error bars represent SEM (8 replicates)). (**b**) The lagging chunk was the major contributor to the Extent of Adaptation (*EoA*) in SM4 but not in SM1 (the error bars represent SEM (8 replicates)). This also means that the contribution of the nose to the *EoA* (which equals (1 – contribution of the lagging chunk)) in SM1 was much more than that of the lagging chunk. (**c) and (d**) Schematic representations of the distribution of efficiency across individuals during adaptation during the initial phases of evolution (before generation 80). Due to the high efficacy of selection in SM1, the majority of individuals were found in the nose (**c**). On the other hand, a relatively low efficacy of selection due to harsher bottlenecks in SM4 resulted in most individuals being found in the lagging chunk (please refer to the text for more details) (**d**). (**e) and (f**) During the later phases of evolution (around generation 360), the contributions of the nose to the overall *EoA* became relatively similar in SM1 and SM4.

The above observations suggest that during the early phases of evolution, populations with higher *g* (here SM4) can face greater impediment (genetic load), which translates into a reduced *EoA*.

### *N_0_/g* is a better predictor of *EoA* than *N_0_g*

Since our simulations suggested that the rate of adaptation is positively related to *N_0_* and negatively related to *g*, we went on to test if *N_0_/g* is a better predictor of adaptive trajectories than *N_0_g*. This turned out to be the case, not only in our simulations (Fig. 6; also see Figs. 2 and S14), but also for our experiments. The *N_0_/g* values of LL, SS and SL populations were approximately 3.01×10^9^, 1.13×10^4^, and 5.02×10^3^, respectively, which led to a predicted *EoA* trend of LL>SS>SL, which was observed for both *K* and *R* in the experiments (Fig. 1).

**Fig. 6.**
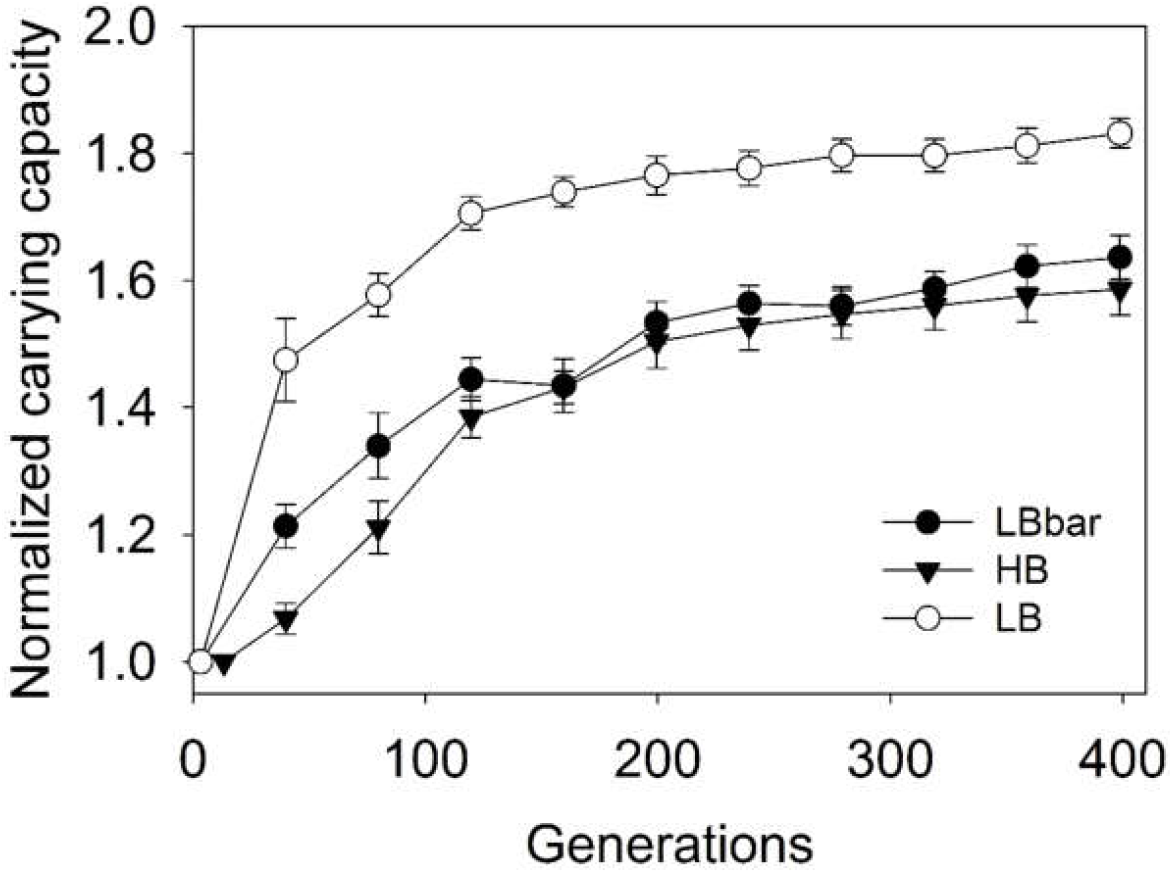
Simulations: *EoA* trajectories in terms of *K*. Populations with similar *N_0_/g* (LBbar and HB) match more closely in terms of mean adaptive trajectories than populations with similar *N_0_g*(LB and HB). LB: *N_0_* ≈ 3600, bottleneck ratio: 1/10; HB: *N_0_* ≈ 900, bottleneck ratio: 1/10^4^; LBbar: *N_0_* ≈ 225, bottleneck ratio: 1/10.

## Discussion

### Overview of the main results and what they suggest

Most experimental studies with periodically bottlenecked asexual populations have used the harmonic mean *(HM)* as the measure of population size (De Visser and Rozen 2005; Desai et al. 2007; Lachapelle et al. 2015; Lenski et al. 1991; Rozen et al. 2008; Samani and Bell 2010; Vogwill et al. 2016) to investigate quantities akin to the extent of adaptation *(EoA)*. Desai et al. (2007) stated that the enhancement in mean population fitness with respect to time (a quantity equivalent to *EoA*) depends upon the harmonic mean of the population size in such populations (Desai et al. 2007). However, there has been no empirical or theoretical test for the validity of *HM* as a predictor of *EoA*. Therefore, as a starting-point, we performed evolutionary experiments on *E. coli* populations to test if the harmonic mean of population size (*HM (*=*N_0_g*)) can predict *EoA*. Our experiments revealed that *N_0_g* does not predict *EoA* (Fig. 1). This observation could be interpreted in two ways. Either there was something wrong with *HM* in terms of predicting *EoA*, or there were some idiosyncratic properties of our experimental system (e.g. different kinds of containers) that masks the relationship between *HM* and *EoA*. Apart from their different numbers (whose effect we study here) and the fact that the LL/SL treatments are grown in flasks while the SS treatment is grown in tissue culture plates, there are no differences between the three treatments, and hence the corresponding selection pressures. Since they are grown with continuous shaking, aeration is unlikely to be a significant issue. To account for the possibility that some idiosyncrasies of our experiments were responsible for our results, and to test if the results of our experimental case-study were generalizable, we simulated the adaptive evolution of asexual populations that grow via fission. For this purpose, we used a very generic Individual-Based Model (IBM) that did not contain any *E. coli* specific functions or parameters. The idea here was that if the outcomes of the simulations matched the experiments, we could be reasonably confident that the experimental results are not due to some peculiarities of the *E. coli* system or experimental protocols. The simulations also revealed no association between *N_0_g* and *EoA* (Fig. 2) which strengthened the first interpretation that *N_0_g* is not a good predictor of *EoA*.

It must be added here that conventionally, *HM* has been treated as an evolutionarily relevant measure of population size only in terms of neutral mutations (Charlesworth 2009; Kimura 1983). However, at least in terms of fixation probabilities of beneficial mutations, it has been shown that population size measures similar to *HM* can act as the relevant measure of population size (Campos and Wahl 2009, 2010; Heffernan and Wahl 2002).

To investigate why *N_0_g* is an inappropriate measure for predicting *EoA*, we used our IBM to test how *EoA* varied with *N_0_* and *g* independently, and found the counter-intuitive result that *EoA* varies negatively with *g* (Figs. 3b and S7). To explain this result, we probed the composition of our simulated populations as they evolved (Figs. 4 and 5). We found that *g* plays a dual role in terms of determining *EoA*. Higher values of *g* positively affect *EoA* by increasing the supply of variation, but negatively affect *EoA* by decreasing the efficacy of selection, as reflected by a consistently greater genetic load. We found that this second effect of *g* on *EoA* overshadows the first, something that is underappreciated in the empirical literature. Since *N_0_* and *g* have positive and negative relationships respectively with *EoA*, intuition suggests that a good predictor of *EoA* should also do the same. One such expression (of the many possible, taking into account that in most evolutionary experiments *N_0_* ≫ *g*) is *N_0_/g*. *N_0_/g* indeed turns out to be a better predictor of *EoA* in our simulations than *N_0_g* (Figs. 6 and S14). We show below how both measures, (i.e. *N_0_g* and *N_0_/g*) could lead to similar predictions about *EoA* under certain circumstances, and why is it important to consider the cases when this correspondence breaks down.

The rest of the discussion elaborates the various insights mentioned above (and some more) and their consequences.

### Periodic bottlenecks lead to increased variation but reduced adaptation

The growth of many natural asexual populations is punctuated by episodic bottlenecks caused by, for example, abrupt dissociation from hosts or spread of infections across hosts (reviewed in (Abel et al. 2015)), etc. Moreover, periodic sampling during sub-culturing is a common feature of most asexual populations propagated during experimental evolution studies (Kawecki et al. 2012; Lenski et al. 1991). Therefore, it is important to appreciate the complex role played by periodic bottlenecks in the evolutionary dynamics of asexual populations.

Most experimental evolution studies with asexual microbes are started with either genetically uniform/clonal replicate populations or a mixed inoculum of relatively small number of genotypes. In such populations, de *novo* beneficial mutations are the principal basis of adaptation (Barrick et al. 2009; Kawecki et al. 2012). That is why populations that experience greater number of binary fissions per generation are expected to generate more *de novo* beneficial variation and thus, to have a higher extent of adaptation (*EoA*). Now, the number of binary fissions per generation is given by *N_0_(2^g^-1)/g*. This quantity varies positively with the number of generations before a bottleneck (*g*) and also with the size of the population at the bottleneck (*N_0_*). Thus, all else being equal, *HM (≈N_0_g)* is expected to be a good predictor of *EoA*.

However, the above line of reasoning disregards the fact that there can be a significant loss of variation during periodic bottlenecks. As *g* increases, *N_0_* represents a smaller fraction of the final population size (*N_f_*) before bottleneck, which in turn increases the chances of loss of variation. For example, assume that there are two bacterial populations that have the same value of *N_0_* (=10^2^) but *g* values of 3.32 and 13.28, leading to *N_f_* values of 10^3^ and 10^6^ respectively (Lenski et al. 1991). For a given value of *N_0_*, increasing the value of *g* decreases the probability that a new beneficial mutation would survive the bottlenecks (Wahl et al. 2002; Wahl and Zhu 2015). All else being equal, this should reduce *EoA*.

Thus, increasing *g* has opposite effects on supply and survival of mutations in a population. Several theoretical studies have investigated which of these two effects is more important for adaptive evolution in asexual populations. For example, it has been suggested that increasing *g* increases the probability of fixation of a beneficial mutation (Heffernan and Wahl 2002). This implies that the positive relationship between *g* and mutational supply can overcome the negative effect of increasing *g* on adaptation. Other theoretical studies have also shown a positive relationship between adaptively relevant population size and the product *N_0_g* (Campos and Wahl 2009, 2010). Unfortunately, this rich body of theoretical predictions are not in the context of quantities (like *EoA*) that are experimentally tractable, which was one of the motivations behind this study.

Our experiments (Fig. 1) and simulations (Fig. 2) showed that populations with similar values of *N_0_g* can have very different adaptive trajectories, suggesting that *N_0_g* is not a good predictor of *EoA*. Moreover, our simulations predicted the relationship between *EoA* and *g* to be negative (Figs. 3b and S7) and not positive. These two results disagree with a rather large body of existing literature, as outlined above. One way by which this can happen is if our model incorporates some atypical assumptions which lead to the observed counter-intuitive results. However, if that were to be the case, then one would also expect our model to show other unintuitive results. Therefore, we first investigated whether various other predictions of our model matched those from the literature.

Our model was able to replicate several intuitive theoretical predictions that had not been coded directly. See Supplementary Data (SD2, also see the associated Figs. S1, S3, S6, and S10). Therefore, it is reasonable to state that our model was generic and a good descriptor of evolving bacterial populations.

### *EoA* varies negatively with *g* because higher *g* makes selection less effective

In order to explain why *EoA* varies negatively with *g*, we simulated populations with similar values of *N*_0_ (*i.e*., bottleneck size) but different degrees of harshness of the bottlenecks, namely SM1 (lenient bottleneck (= 1/10), *g* =3.32) and SM4 (harsh bottleneck (= 1/10^4^), *g* = 13.28) (Figs. 5 and 6).

Our results demonstrate that higher *g* decreases the efficacy of selection in terms of spreading beneficial mutations and purging deleterious ones (Fig. 5, also see Figs. S11 and S12). As shown in Fig. 5, very high-efficiency classes rise to very high frequencies in SM1 populations by generation 80. However, such classes fail to do so in SM4 populations. Owing to lenient bottlenecks (lower *g*), selection operates so effectively in SM1 that its best efficiency class quickly converges with the modal class (Fig. S11b). This is also reflected by the proximity of the mean class with the modal class in SM1 (Fig. S11c). Thus, once a high-fitness class arises in an SM1 population, its rapid spread results in a steep increase in the population’s *EoA*. However, despite having the same bottleneck size (= *N_0_*) as SM1, SM4 populations exhibit a much slower rise in their *EoA*. This happens due to two reasons. As opposed to SM1, high-fitness genotypes in SM4 need to rise to much higher frequencies to survive the harsh periodic bottlenecks. This results in the removal (due to sampling) of several high-fitness classes from SM4 during the bottleneck (Fig. S12a). More importantly, the higher mutational supply rate of SM4 increases the genetic load (Fig. 5), which ultimately results in a much slower rise in the *EoA* of SM4.

### Evolution of carrying capacity can feedback into adaptive trajectories

Both our experiments and simulations showed that carrying capacity (*K*) can evolve during adaptation in asexual microbes (Figs. 1a and 2a respectively), which is consistent with previous results (Novak et al. 2006). Unfortunately, most models of asexual adaptation do not take into account such adaptive changes in the carrying capacity (Campos and Wahl 2009, 2010; Gerrish and Lenski 1998; Wahl and Gerrish 2001). Most evolution experiments keep the bottleneck ratio (represented by *g*) constant (Kawecki et al. 2012; Lenski et al. 1991). This constancy of *g* ensures that any evolutionary change in carrying capacity would also change *N_0_*. In other words, if *K* increases, a constant value of *g* throughout evolution would ensure an increase in *N_0_*. Since higher values of *N_0_* accelerate adaptation (Fig. 3a), the regularity of bottlenecks introduces a positive feedback during evolution if *K* increases adaptively. Stated differently, a larger value of *N_0_* would make a population evolve higher K, which in turn would increase the next *N_0_*, and so on. We think that this aspect of fitness should not be omitted from theoretical models of how microbes evolve, particularly under resource-limited conditions, which are a common feature of experimental evolution protocols (Kawecki et al. 2012; Lenski et al. 1991).

### *N_0_/g* is a better predictor of *EoA* than *N_0_g*

As shown in Figs. 3b, S7 and 4, when selection is at work, *EoA* decreases with increasing *g*. This suggests that a population size measure which is an increasing function of *N_0_* but a decreasing function of *g* can be a better predictor of *EoA* than the conventional formula (*HM*). For example, as shown in Fig. 6, we found that *N_0_/g* is a better predictor of *EoA* than *HM* (= *N_0_g*). Admittedly, it is not possible to reason from this that the expression *N_0_/g* will always be a good predictor of *EoA*, and we make no such claims. We simply submit this expression as a potential candidate for this purpose and hope that future theoretical work will be able to validate this empirically derived quantity.

Most theoretical studies assume that the final population size attained in their study systems (*N_f_*) is constant (Campos and Wahl 2009, 2010; Gerrish and Lenski 1998; Heffernan and Wahl 2002; Wahl et al. 2002; Wahl and Gerrish 2001). Interestingly, if the experimental populations that are being compared have similar values of *N_f_* (Desai et al. 2007; Y. Raynes et al. 2014; Vogwill et al. 2016), then the populations with larger values of *N_0_g* will typically also have larger values of any quantity that is an increasing function of *N_0_* but a decreasing function of *g*. This is because of two reasons. First, if *N_f_* is held constant, since *N_f_* = *N_0_2^g^*, increasing *N_0_* necessarily decreases *g*. Second, in most empirical studies, *N_0_≫g*. Consequently, if *N_f_* is assumed to be the same the populations being compared, any prediction based on the relative values of *N_0_g* will typically be similar to predictions based on *N_0_/g* (Fig. S15). However, whenever *N_f_* is not held constant (e.g., Figs. 2, 3, 4, 6 and S14, and studies like (Lachapelle et al. 2015; Ramsayer et al. 2013; Y. Raynes et al. 2014; Rozen et al. 2002; Samani and Bell 2010), *N_0_/g* predicts *EoA* much better than *N_0_g*. The above observations can explain why *N_0_g* has been widely used across several empirical studies despite failing to capture the effects of *g* on *EoA* accurately.

### Implications of our results

At very long time-scales, the high-fitness mutations accessible only to SM4 (but not to SM1) may end up surviving a harsh periodic bottleneck. A post-facto analysis of our SM4 simulations shows that mutations of this kind rise to a frequency between 10 ^−7^ and 10^−8^ in a typical growth phase just prior to bottlenecks in SM4. Since *N_0_* is close to 10^3^ in these populations, the above high-quality mutations would survive one bottleneck in every 10^4^ to 10^5^ growth phases which roughly amounts to 1.3×10^5^ to 1.3×10^6^ generations. However, to this date, there are no reported experimental evolution studies over this long a time-span. Therefore, we conclude that the observation that increasing *g* decreases *EoA* should be relevant for the time-scales most commonly employed in experimental evolution studies.

Our results can be used to compare the extents of adaptation in independent evolution experiments with similar environments but dissimilar demographic properties (differences in terms of *N_0_* and/or *g* and/or *N_f_*). Such studies, which compare populations evolving in similar environments but with dissimilar demographic properties, are reasonably common in the field of experimental evolution (Desai et al. 2007; Lachapelle et al. 2015; Y. Raynes et al. 2014, 2014; Rozen et al. 2002; Samani and Bell 2010; Vogwill et al. 2016).

Our study shows that in serially bottle-necked asexual populations, the destructive aspect of bottlenecks (reduction in efficacy of selection by harsher bottlenecks) can overshadow their constructive aspect (increase in supply of variation in harsher bottlenecks). This calls for a change in perspective about periodic bottlenecks and a substantial re-evaluation of the role of population size as a predictor of adaptive evolution.

## Supplementary information

### Supplementary Methods

#### Experimental Evolution

##### Selection regimens

We propagated three distinct regimens (LL, SL, and SS) of *Escherichia coli* MG 1655 populations for more than 380 generations in Nutrient Broth with a fixed concentration of an antibiotic cocktail containing a mixture of three antibiotics at sub-lethal concentrations. 8 independently evolving replicate lines each of LL (<u>l</u>arge *HM* and <u>l</u>arge *N_f_*, culture volume: 100ml (flasks)), SL (<u>s</u>mall *HM* but large *N_f_*, culture volume: 100ml (flasks)), and SS (<u>s</u>mall *HM* and <u>s</u>mall *N_f_*, culture volume: 1.5 ml (well plates)) were derived from a single *Escherichia coli* K-12 MG1655 colony and propagated during the course of our evolution-experiment. It is possible that the differences in culture containers or absolute numbers at the stationary phase could have exposed these populations to somewhat different selection pressures. However, our simulations had no such differences coded into them and yet yielded identical results. Thus, it is highly unlikely that the empirically observed differences between the SS, SL and LL populations arose due to differences in culture conditions. All the cultures were propagated at 37°C, and were shaken continuously at 150 rpm. The three population regimens experienced different numbers of evolutionary generations (*g*) between periodic bottlenecks (*i.e*., before they were sub-cultured). SS and SL had similar *HM (i.e*., *N_0_g*) albeit obtained through different combinations of *N_0_* (SS>SL) and *g* (SL>SS) such that the *N_f_* of SL was approximately 73 times larger than that of SS. The *N_f_* of SL was similar to that of LL, while the harmonic mean size of LL was > 16,500 times larger than that of SL and SS (Table S1). The bottleneck ratio for random sampling at the end of each growth phase was 1/10 for LL, 1/10^4^ for SS, and 1/10^6^ for SL. 1 ml cryostocks belonging to each of the twenty four independently evolving populations were stored periodically. This scheme automatically gave rise to the following rank-distribution in terms of the quantity *N_0_/g*: SL<SS<LL.

#### Culture environment for experimental evolution

24 independent bacterial populations *Escherichia coli* K-12 MG1655 were grown in Nutrient Broth with a fixed concentration of an antibiotic cocktail containing a mixture of three antibiotics at sub-lethal concentrations:

1. Norfloxacin (0.015 μg/ml)
2. Rifampicin (6 μg/ml)
3. Streptomycin (0.1 μg/ml)

Composition of Nutrient Broth (Himedia Laboratories Pvt. Ltd.):

- Peptic digest of animal tissue (5 g/l)
- Sodium chloride (5 g/l)
- Beef extract (1.50 g/l)
- Yeast extract (1.50 g/l)

The following table summarizes the numerical properties of the three population regimes used in our study:

**Table S1.**
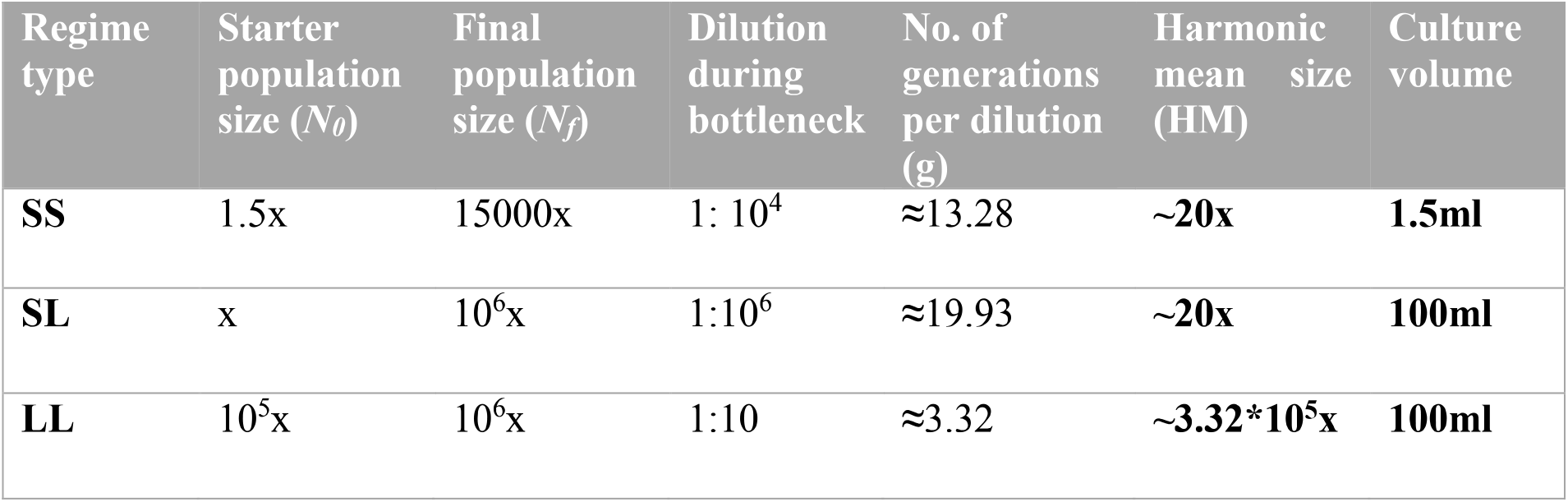
A summary of the experimental populations. x ≈ 10^5^ in our experiments.

##### Fitness assays

To reconstruct the evolutionary trajectories of our experimental bacterial populations, we measured bacterial growth using an automated multi-well plate reader (Synergy HT, BIOTEK ^®^ Winooski, VT, USA). Bacterial growth was measured in the same medium (containing the antibiotic cocktail) that the populations experienced during evolution. We used OD at 600 nm as a proxy for bacterial growth. Bacteria from the cryostocks belonging to each of the 24 populations were grown in 96 well plates. Each cryostock-derived population was assayed in three measurement-replicate wells in a 96 well plate. Each well contained 180 μl growth medium (Nutrient Broth with the antibiotic cocktail) containing 1:10^4^ diluted cryostock. The plate was incubated at 37°C, and shaken continuously by the plate-reader throughout the growth assay. OD readings taken every 20 minutes during this incubation resulted in sigmoidal growth curves. Fitness measurements were done using cryostocks belonging to multiple time-points in order to reconstruct evolutionary trajectories. While reconstructing fitness trajectories, it was made sure that every 96 well-plate contained populations belonging to similar time-points (in terms of number of generations) during the course of evolution. We used the carrying capacity (*K*) and maximum population-wide growth rate (*R*) as the measure of fitness (Novak et al. 2006)). *K* of a population was defined as the maximum OD value attained over a period of twenty four hours (the highest value in the sigmoidal growth curve) (Karve et al. 2016; Novak et al. 2006). *R* was estimated as the maximum slope of the growth curve over a running window of four OD readings (each window spanning one hour) (Karve et al. 2015, 2016; Vogwill et al. 2016). Fitness measurements were done using cryostocks belonging to multiple time-points in order to reconstruct evolutionary trajectories.

#### Statistics

Repeated measures ANOVA were performed independently for each of the two growth parameters (*K* and *R*). “Regimen-type” (LL/SL/SS) was treated as the categorical factor, and TIME (9 time-points) as the repeated measures factor. We also include the interaction of the two factors (Regimen-type and TIME) in the ANOVA model. We did not perform post-hoc tests in the RM ANOVA analysis as it involved very large number of comparisons that were irrelevant for the present question and thus reduced statistical power. Instead, we analyzed the two growth parameters (*K* and *R)* independently at each time point in the *EoA* trajectory (Fig 1A and 1B) using nested-design ANOVAs with population regimen-type (SS, SL or LL) as the fixed factor and replicate-line (1-8, nested in population-type) as the random factor. For each of these ANOVAs, we further corrected the *p*-value of the main-effect of “Regimen-type” using the Holm-Šidàk correction (Abdi 2010) to control the family-wise error rate. The means from all those ANOVAs which showed a significant (< 0.05) *p*-value after the Holm-Šidàk correction, were further subjected to Tukey’s HSD to identify which pair-wise differences were significant.

### Algorithm for the individual based model used in this study

Our model simulates the growth of individual bacteria in density-dependent resource-limited conditions. Each bacterium is represented by an array which has the following components:

1. Determinant of efficiency (K_eff): determines how much food can be assimilated per unit time
2. Determinant for threshold (thres): how much food needs to be assimilated in order to divide
3. Bodymass: how big is the bacterium (where is it along its cell cycle) at any given time

At the beginning of the simulation, two global scaling quantities, Food_Proxy and Body_Proxy are declared for the whole population. As the names suggest, Food_Proxy acts as a proxy for the amount of available resources initially, while Body_Proxy (=250) is a proxy for bodymass of the ancestor. Each simulation run is started with 100 individuals and each individual is allotted K_eff_i_ value given as Eff_i_ * Food_Proxy. Here, Eff_i_ is a random number picked from a uniform distribution U(0.95-1.05) and K_eff_i_ determines how much food would be consumed in a density–dependent manner and when its food consumption would stop (as per the conditions given below). Similarly, the parameter for the threshold for each of the 100 starting individuals is assigned as a random number picked from a uniform distribution between 0.95*(Body_Proxy) and 1.05*(Body_Proxy).

Each bacterium has the same initial biomass (an arbitrarily small quantity, 10 units in this case).

Time is implicitly defined in our code and each iteration signifies one unit of time.

In each iteration, each bacterium “grows and divides” according to the following rules:

*If for bacterium i, (population size/ K_eff_i_) ≥ 1, it doesn’t eat anything: its bodymass remains the same as the earlier iteration*.
*If for bacterium i (population size/ K_eff_i_) < 1, it eats 10*(1 - (population size/ K_eff_i_)) units of food in this iteration: its bodymass increases by 10*(1- (population size/K_eff_i_) units)*.
*If at the end of this iteration, bodymass_i_ > thres_i_, bacterium i divides into two equal parts. A small thermodynamic cost (constant for all individuals) is deducted so that the sum of the bodymass of the daughter cells is exactly 1 unit less than the bodymass of the mother cell at the time of division*.
*If the bacterium divides, there is a 1 in 100 chance for each of the daughter cells that it mutates. If a mutation occurs, the new parameter for efficiency is drawn from an already defined normal distribution that is used throughout the simulation. The same applies to threshold. (Threshold and efficiency mutate independently in each bacterium.)*
*The total size of the population is saved at the end of every iteration. The total amount of food consumed during each iteration is also computed*.

The above description (italics) represents all the processes that happen within an iteration.

The process is repeated (and the population grows) until the following conditions are fulfilled:

1. The number of iterations is greater than 2000.
2. The amount of food consumed during each iteration < 0.08* Food_Proxy

If the above conditions are met simultaneously, food consumption is stopped, a defined fraction of individuals are sampled randomly and the whole process is started with this sample population (this represents bottlenecking). The above process is continued for *q* bottlenecks. The bottleneck ratio and the number *q* are predefined, depending upon the type of population being studied. This gives rise to *q* sigmoidal growth curves. Two quantities are extracted from each sigmoidal curve:

1. Carrying capacity (*K*, the maximum size of the population in each growth phase)
2. Maximum linear growth rate (*R*, the maximum slope of population growth over 100 iterations). Straight lines were fit on overlapping moving windows of 100 iterations on the entire time-series of population size values within each growth-phase. The maximum value of the slope observed within the entire time-series of population size values (sigmoidal curve) was taken to be the maximum growth rate (*R*).

Time series of carrying capacities and maximum growth rates are computed using the series of *q* sigmoidal curves.

In each simulation used in this study, the carrying capacity of the first growth phase was ≈ 1.8*Food_Proxy. The value of Food_Proxy was adjusted in such a way that it gave rise to the desired value of the carrying capacity of the first growth phase in a simulation. The carrying capacities corresponding to the subsequent growth phases was an emergent result of Darwinian evolution in the simulations. A summary of the key differences between our model and previous theoretical studies is given in Table S2.

**Table S2.**
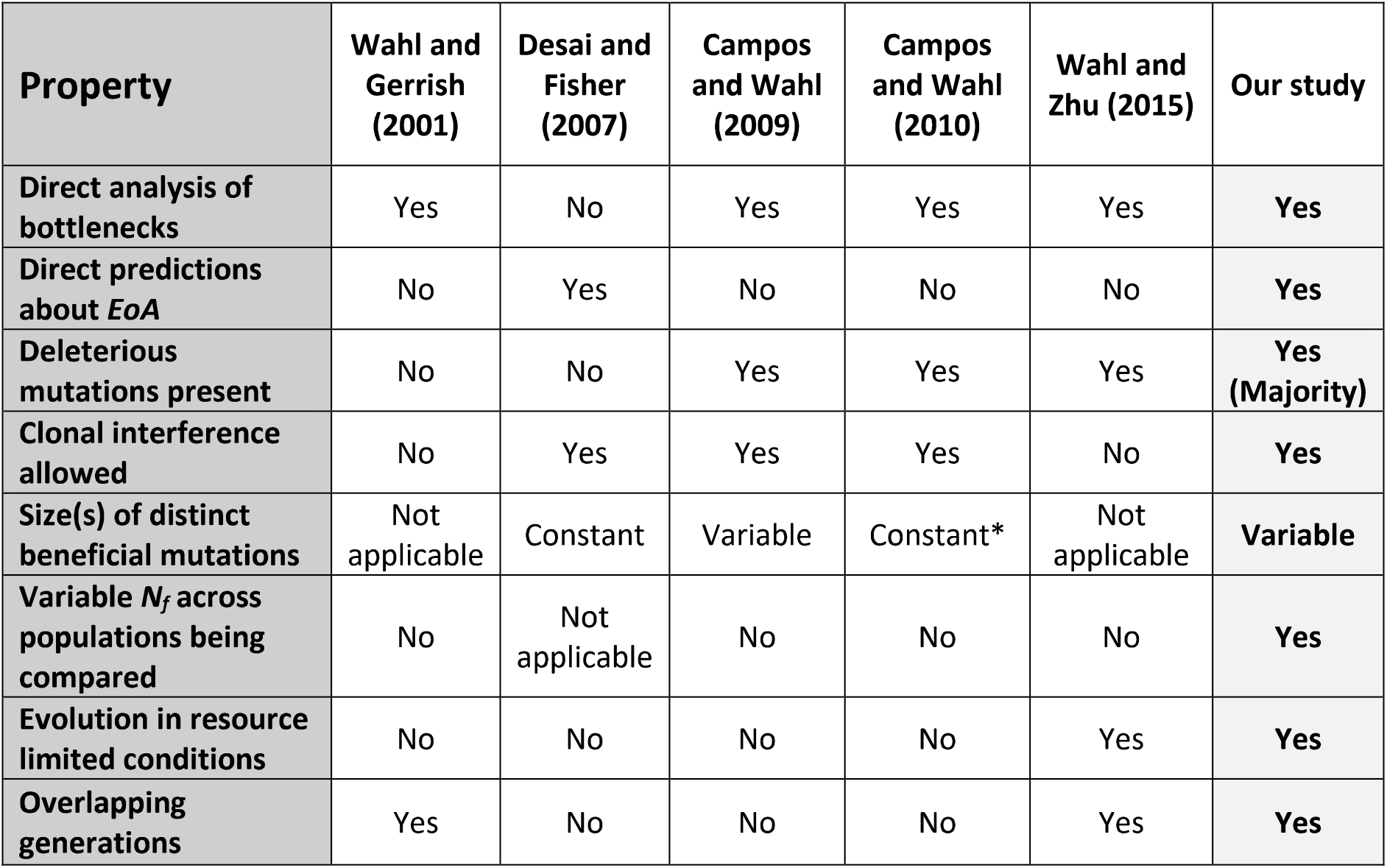
A summary of the key differences between our model and previous theoretical studies.

| Property | Wahl and Gerrish (2001) | Desai and Fisher (2007) | Campos and Wahl (2009) | Campos and Wahl (2010) | Wahl and Zhu (2015) | Our study |
| --- | --- | --- | --- | --- | --- | --- |
| Direct analysis of bottlenecks | Yes | No | Yes | Yes | Yes | Yes |
| Direct predictions about *EoA* | No | Yes | No | No | No | Yes |
| Deleterious mutations present | No | No | Yes | Yes | Yes | Yes (Majority) |
| Clonal interference allowed | No | Yes | Yes | Yes | No | Yes |
| Size(s) of distinct beneficial mutations | Not applicable | Constant | Variable | Constant* | Not applicable | Variable |
| Variable $N_f$ across populations being compared | No | Not applicable | No | No | No | Yes |
| Evolution in resource limited conditions | No | No | No | No | Yes | Yes |
| Overlapping generations | Yes | No | No | No | Yes | Yes |

*While assuming that distinct beneficial mutations can have different effects, Campos and Wahl (2010) arrive at an expression of *N_e_* that itself depends upon the size of such effects (***s_b_***). They then state that “Clearly an effective population size that depends on ***s_b_*** is unsatisfactory.” Then they assume that “all beneficial mutations have the same effect” to arrive at a much simpler expression.

The following simulation settings were used in our study:

| Population type | Food_proxy | Bottleneck ratio | Number of bottlenecks (400 generations) |
| --- | --- | --- | --- |
| XX' | $2 \times 10^4$ | 1/10 | 120 |
| SS' | $10^5$ | 1/10<sup>2</sup> | 60 |
| SL' | $5 \times 10^6$ | 1/10<sup>4</sup> | 30 |
| BN1 | $5 \times 10^3$ | 1/10 | 120 |
| BN2 | $5 \times 10^4$ | 1/10<sup>2</sup> | 60 |
| BN3 | $5 \times 10^5$ | 1/10<sup>3</sup> | 40 |
| BN4 | $5 \times 10^6$ | 1/10<sup>4</sup> | 30 |
| SM1 | $5 \times 10^3$ | 1/10 | 120 |
| SM4 | $5 \times 10^6$ | 1/10<sup>4</sup> | 30 |
| HB | $5 \times 10^6$ | 1/10<sup>4</sup> | 30 |
| LB | $2 \times 10^4$ | 1/10 | 120 |

| Population type | Food_proxy | Bottleneck ratio | Number of bottlenecks<br>(400 generations) |
| --- | --- | --- | --- |
| LBbar | $1.25 \cdot 10^3$ | 1/10 | 120 |
| MBbar | $2.5 \cdot 10^4$ | 1/10<sup>2</sup> | 60 |
| SNFBN1 | $5 \cdot 10^4$ | 1/10 | 120 |
| SNFBN2 | $5 \cdot 10^4$ | 1/10<sup>2</sup> | 60 |
| SNFBN4 | $5 \cdot 10^4$ | 1/10<sup>4</sup> | 30 |
| Fig. S6a (small<br>bottleneck size) | $5 \cdot 10^4$ | 1/10 | 120 |
| Fig. S6a (large<br>bottleneck size) | $5 \cdot 10^6$ | 1/10 | 120 |
| Fig. S6b (small<br>bottleneck size) | $5 \cdot 10^4$ | 1/10<sup>3</sup> | 40 |
| Fig. S6B (large<br>bottleneck size) | $5 \cdot 10^6$ | 1/10<sup>3</sup> | 40 |
| Fig. S6c (small<br>bottleneck size) | $5 \cdot 10^5$ | 1/10<sup>4</sup> | 30 |
| Fig. S6c (large<br>bottleneck size) | $5 \cdot 10^6$ | 1/10<sup>4</sup> | 30 |
| Fig. S7a (lenient<br>bottleneck) | $5 \cdot 10^2$ | 1/10 | 120 |
| Fig. S7a (harsh<br>bottleneck) | $5 \cdot 10^6$ | 1/10<sup>5</sup> | 24 |
| Fig. S7b (lenient<br>bottleneck) | $5 \cdot 10^4$ | 1/10 | 120 |
| Fig. S7b (harsh<br>bottleneck) | $5 \cdot 10^6$ | 1/10<sup>3</sup> | 40 |
| Fig. S7c (lenient<br>bottleneck) | $5 \cdot 10^5$ | 1/10 | 120 |
| Fig. S7c (harsh<br>bottleneck) | $5 \cdot 10^7$ | 1/10<sup>3</sup> | 40 |
| Fig. S7d (lenient<br>bottleneck) | $5 \cdot 10^6$ | 1/10 | 120 |
| Fig. S7c (harsh<br>bottleneck) | $5 \cdot 10^7$ | 1/10<sup>2</sup> | 60 |

We checked if our simulations met several other theoretical expectations from the extant literature. As expected, despite following the same distribution for mutations, large populations showed curves of diminishing returns while adapting, whereas very small populations showed stepwise increase in fitness with long periods of stasis (Fig. S1). This happens because very small populations (but not large ones) need to wait for beneficial mutations to arise (Sniegowski and Gerrish 2010). Moreover, the extent of adaptation showed a positive but saturating relationship with an unambiguous measure of absolute population size in our simulations (Fig. 3a and S7a). The simulations also revealed a non-monotonous relationship between fitness and mutation rate (Orr 2000) (Fig. S7).

**Table S3.**
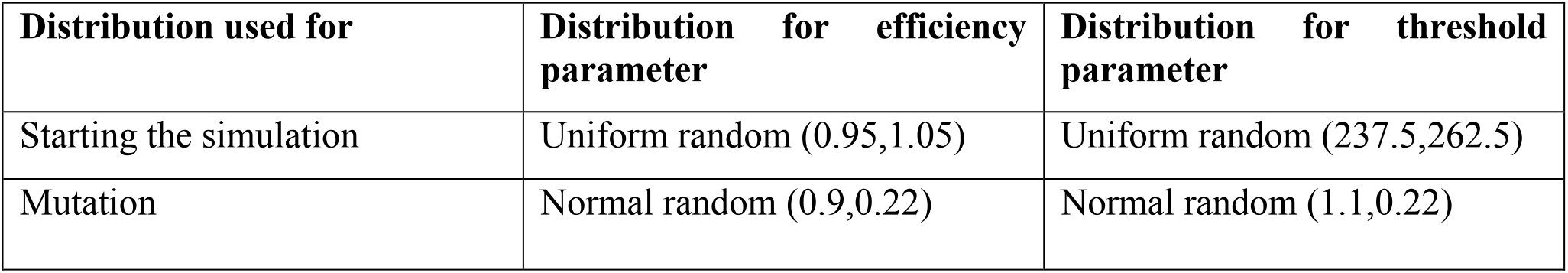
Distributions for parameters used in simulations.

#### Supplementary Data

##### SD1. Statistical analysis of empirical results

**Table S4.**
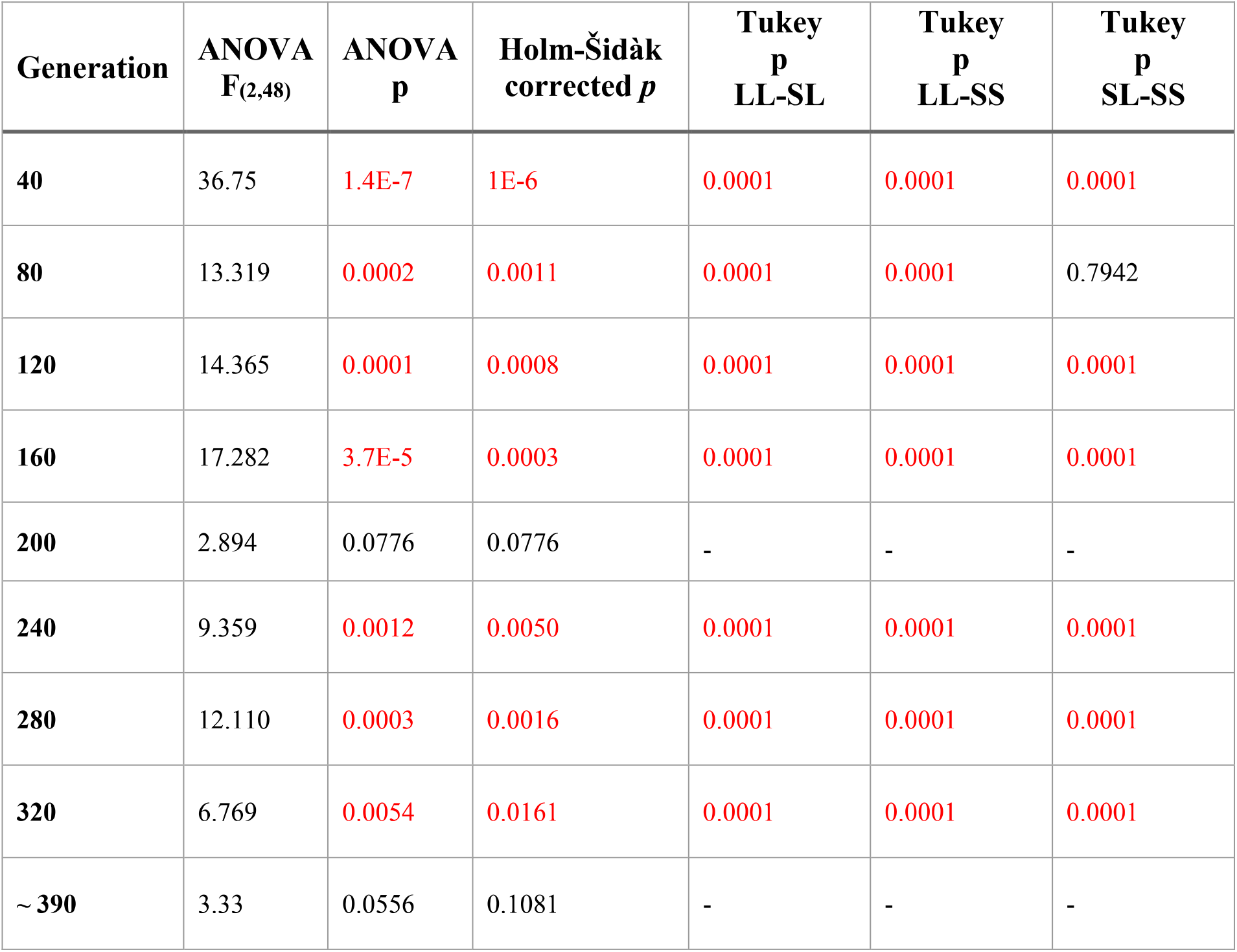
A summary of statistical analysis of carrying capacity (*K*) measurements in empirical populations. The values in red represent statistically significant difference (*p*<0.05). The *p*-values corresponding to nine independent ANOVAs (corresponding to nine different time points) were subjected to Holm-Šidàk correction. Post-hoc (Tukey) comparisons were done only in cases where the ANOVA *p*-values were less than 0.05 after Holm-Šidàk correction. These post–hoc comparisons were done for three pairwise differences (LL-SL, LL-SS, and SL-SS) at each time point. Holm-Šidàk correction was not done on Tukey *p*-values. The *p*-values are reported to four decimal places.

| Generation | ANOVA<br>$F_{(2,48)}$ | ANOVA<br>$p$ | Holm-Šidák<br>corrected $p$ | Tukey<br>$p$<br>LL-SL | Tukey<br>$p$<br>LL-SS | Tukey<br>$p$<br>SL-SS |
| --- | --- | --- | --- | --- | --- | --- |
| 40 | 36.75 | 1.4E-7 | 1E-6 | 0.0001 | 0.0001 | 0.0001 |
| 80 | 13.319 | 0.0002 | 0.0011 | 0.0001 | 0.0001 | 0.7942 |
| 120 | 14.365 | 0.0001 | 0.0008 | 0.0001 | 0.0001 | 0.0001 |
| 160 | 17.282 | 3.7E-5 | 0.0003 | 0.0001 | 0.0001 | 0.0001 |
| 200 | 2.894 | 0.0776 | 0.0776 | - | - | - |
| 240 | 9.359 | 0.0012 | 0.0050 | 0.0001 | 0.0001 | 0.0001 |
| 280 | 12.110 | 0.0003 | 0.0016 | 0.0001 | 0.0001 | 0.0001 |
| 320 | 6.769 | 0.0054 | 0.0161 | 0.0001 | 0.0001 | 0.0001 |
| ~ 390 | 3.33 | 0.0556 | 0.1081 | - | - | - |

**Table S5.** A summary of statistical analysis of maximum growth rate (*R*) measurements in empirical populations. The values in red represent statistically significant difference (*p*<0.05). The values in red represent statistically significant difference (*p*<0.05). The *p*-values corresponding to nine independent ANOVAs (corresponding to nine different time points) were subjected to Holm-Šidàk correction. Post-hoc (Tukey) comparisons were done only in cases where the ANOVA *p*-value was less than 0.05 after Holm-Šidàk correction. These post–hoc comparisons were done for three pairwise differences (LL-SL, LL-SS, and SL-SS) at each time point. Holm-Šidàk correction was not done on Tukey *p*-values. The *p*-values are reported to four decimal places.

| Generation | ANOVA<br>F<sub>(2,48)</sub> | ANOVA<br>p | Holm-Šidák<br>corrected p | Tukey<br>p<br>LL-SL | Tukey<br>p<br>LL-SS | Tukey<br>p<br>SL-SS |
| --- | --- | --- | --- | --- | --- | --- |
| 40 | 56.631 | 3.5E-9 | 3.1E-8 | 0.0001 | 0.0001 | 0.0001 |
| 80 | 5.996 | 0.0087 | 0.0344 | 0.0001 | 0.0001 | 0.3229 |
| 120 | 12.027 | 0.0003 | 0.0023 | 0.0001 | 0.0001 | 0.0001 |
| 160 | 12.287 | 0.0003 | 0.0023 | 0.0001 | 0.0001 | 0.0001 |
| 200 | 3.093 | 0.0665 | 0.1286 | - | - | - |
| 240 | 8.757 | 0.0017 | 0.0103 | 0.0001 | 0.0001 | 0.0001 |
| 280 | 7.785 | 0.0030 | 0.0147 | 0.0001 | 0.0001 | 0.0001 |
| 320 | 4.096 | 0.0315 | 0.0915 | - | - | - |
| ~ 390 | 0.925 | 0.4122 | 0.4122 | - | - | - |

###### Effect sizes for the differences in *EoA* of SL and SS in terms of *K* and *R*

**Table S6.** Analysis of the differences in *EoA* of populations with similar *HM* in terms of effect sizes. Cohen’s d (Cohen 1988) was used to determine the effect sizes of the differences in the *EoA* of SS and SL. 0.2 < d < 0.5 was interpreted as small effect, 0.5 < d < 0.8 as medium effect, and d > 0.8 as large effect. The majority of differences between SL and SS were found to be of large effect size. The analysis was performed only at time points when the post-hoc (Tukey) p – values corresponding to SL-SS were < 0.05.

| Fitness measure | Generation | Cohen's *d* | Inference about Effect Size |
| --- | --- | --- | --- |
| ***K*** | 40 | 1.112 | Large effect |
|  | 120 | 0.527 | Medium effect |
|  | 160 | 0.786 | Medium effect |
|  | 240 | 0.606 | Medium effect |
|  | 280 | 0.514 | Medium effect |
|  | 320 | 1.000 | Large effect |
| ***R*** | 40 | 1.172 | Large effect |
|  | 120 | 0.770 | Medium effect |
|  | 160 | 1.024 | Large effect |
|  | 240 | 1.037 | Large effect |
|  | 280 | 0.836 | Large effect |

##### SD2. Results from individual-based simulations that match intuitive theoretical predictions that had not been coded directly

Firstly, as expected (Elena et al. 1996; Sniegowski and Gerrish 2010), very small populations showed discontinuous staircase-like (stepwise) trajectories of fitness increase whereas large populations showed smooth adaptive trajectories (Fig. S1). Secondly, *EoA* trajectories showed diminishing returns with time despite never hitting the explicitly coded wall of adaptive limit (Figs. 2, 3, 4, 5, S2a, S3, S4, S6, S7, and S8) (Lenski et al. 1991; Tenaillon et al. 2016). Thirdly, as expected, we found a non-monotonous relationship between *EoA* and mutation rate (Fig. S8c) (Orr 2000). Fourthly, *EoA* showed a positive but saturating relationship with *N_0_* (which is an unambiguous measure of absolute population size) (Fig. 3a) (Gerrish and Lenski 1998; Sniegowski and Gerrish 2010). All this was highly unlikely if our model incorporated unrealistic or atypical assumptions. Furthermore, for numerically similar populations (i.e. populations with similar *N_0_* and *g*) and identical time-frames, the results from our simulations were a very close match to the results of our experiments in terms of both *K* and *R* trajectories (Fig. S3). This again suggests that the IBM captures at least some features of the *EoA* of our *E. coli* populations. Finally, the *EoA* rank predictions generated for the three experimental populations based on our model agreed well with the empirical data (LL>SS>SL, Fig. 1).

##### Qualitative differences in *EoA* trajectory-shapes brought about by large differences in population size

**Fig. S1.**
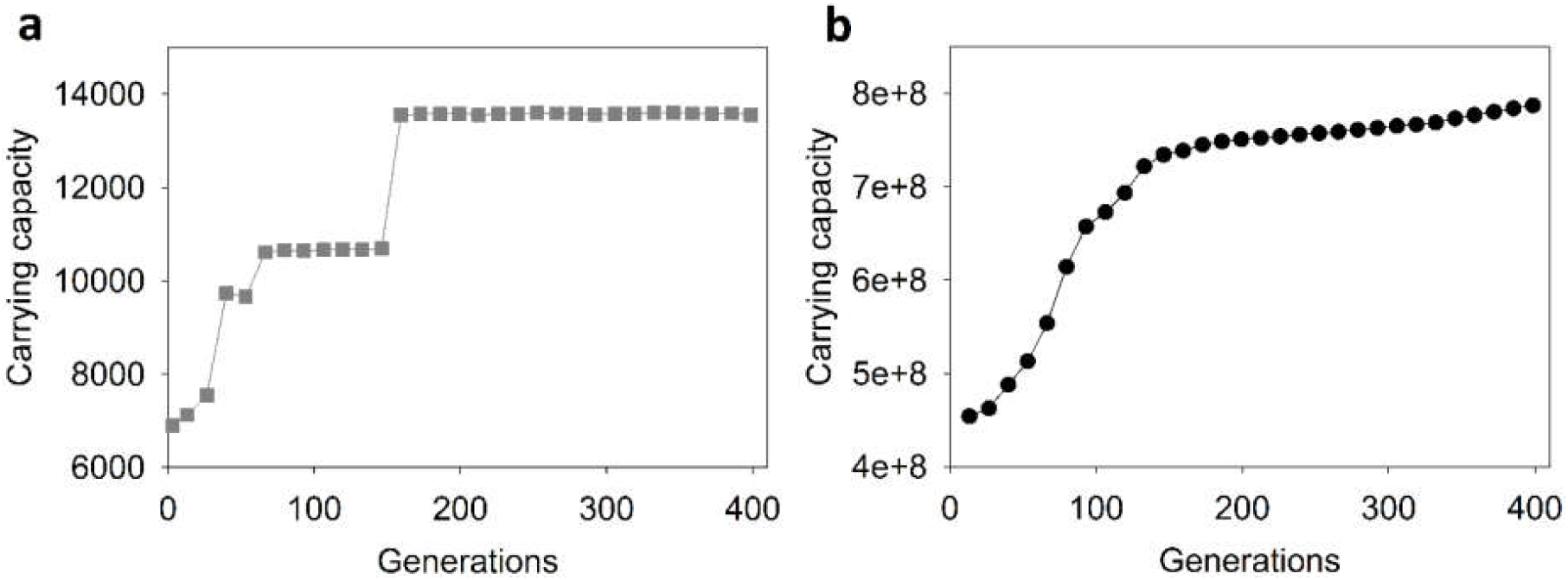
Qualitative differences in adaptive trajectories corresponding to populations with a large different in their sizes. Stepwise increase in fitness (with long periods of stasis) occurred in typically small populations such as the one shown in (**a**) as compared to smooth curves of diminishing returns in typically large populations such as the one shown in _(b)_ (See the ordinates for absolute ranges of *N_f_* during adaptation). The population shown in (**a**) experienced a periodic bottleneck of 1/10 while he population shown in (**b**) was bottlenecked 1/10^4^ periodically.

As expected (Sniegowski and Gerrish 2010), very small populations showed staicase-like (stepwise) trajectories of fitness increase.

##### Changes in standard deviation with sample size

**Fig. S2.**
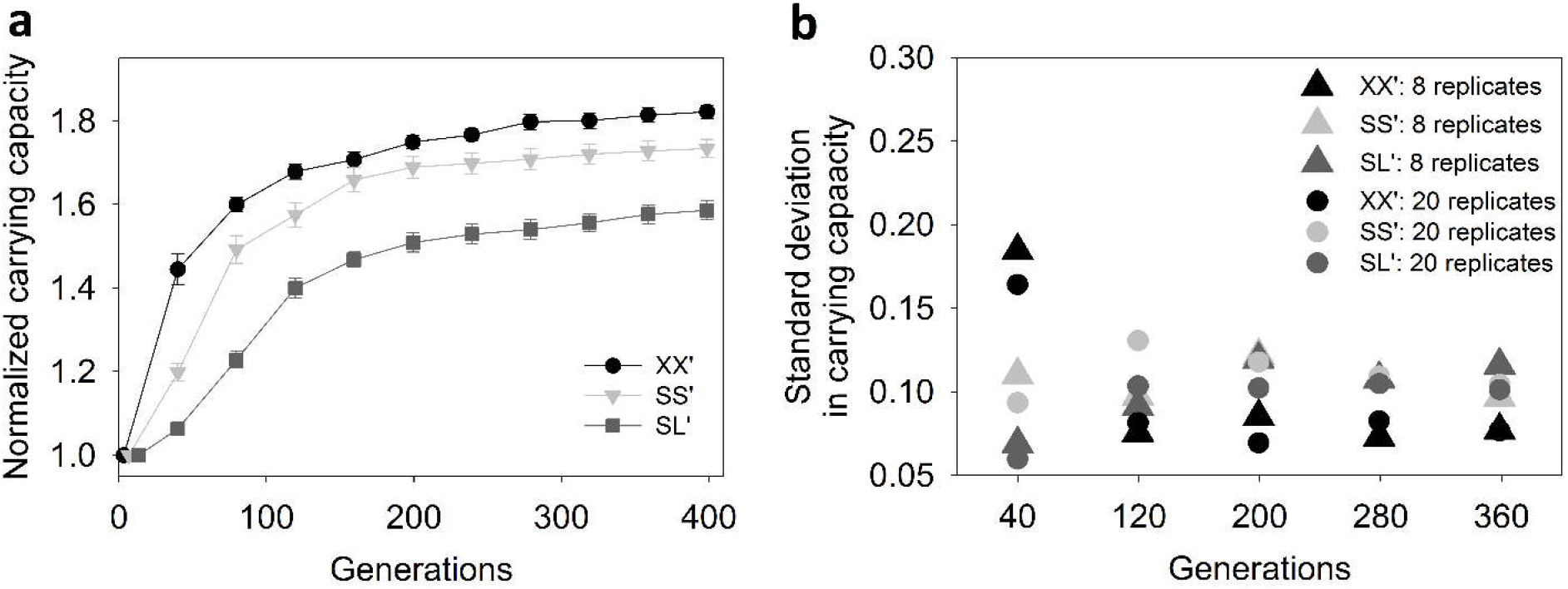
Increasing the number of replicate simulations from 8 to 20 did not result in increase in variation across replicates. (**a**) Increasing the number of independent replicate simulations from 8 to 20 didn’t result in qualitative changes (ranks) of three populations with similar harmonic mean size but different N_0_/g (mean ± SEM; N=20) (compare with Fig. 2a in the Main-text). (**b**) This increase in replicate number also didn’t result in major changes in the standard deviation in carrying capacity during the course of adaptation. XX’: N_0_ ≈ 3.6*10^3^, bottleneck ratio: 1/10; SS’: *N*_0_ ≈ 1.8*10^3^, bottleneck ratio: 1/10^2^; SL’: *N*_0_ ≈ 9*10^2^, bottleneck ratio: 1/10^4^.

Since our simulations are agent-based (and consequently take a very long time to run), we decided to operate on a sample size of 8 replicates per population type throughout our study.

##### Agreement between experiments and simulations

**Fig. S3.**
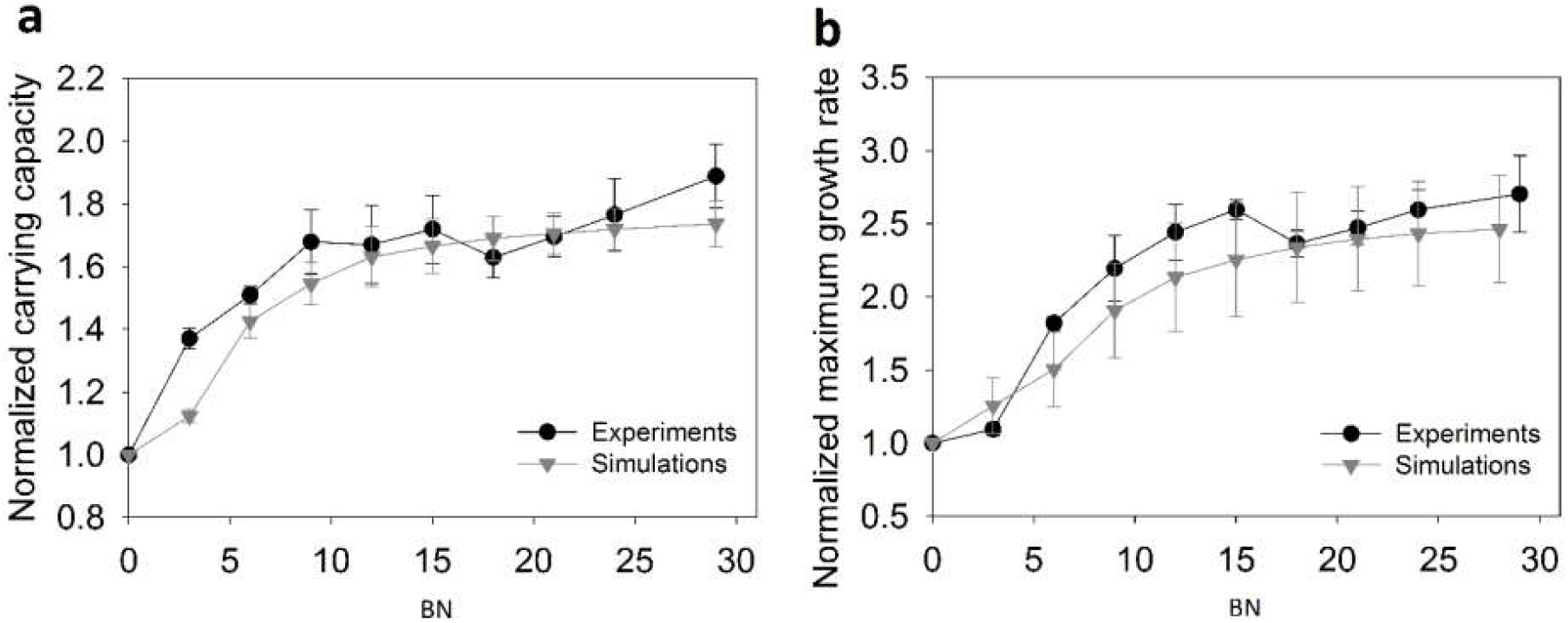
Agreement between experiments and simulations in terms of adaptive dynamics over identical time-scales in numerically similar populations. (**a**) Carrying capacity *(K)* versus bottleneck number *(BN)* (**b**) Maximum growth rate *(R)* versus bottleneck number *(BN)*. Data points represent mean ± SD over 8 replicates. Each data point corresponds to the respective measure of fitness *(K or R)* derived from the sample taken after *BN* bottlenecks. Range of population size: *N_0_* ≈ 10^4.5^; *N_f_* ≈ 10^8.5^; bottleneck ratio = 1/10^4^. Each bottleneck corresponds to approximately 13.28 generations. Both carrying capacity and maximum growth rate are normalized with the ancestral values.

The results of our experiments and simulations agree well in terms of the range and dynamics of adaptation over identical time-scales in numerically similar populations. This applies to both measures of population-level fitness: carrying capacity *(K)* and maximum growth rate *(R)*.

##### Adaptation in terms of efficiency and threshold

**Fig. S4.**
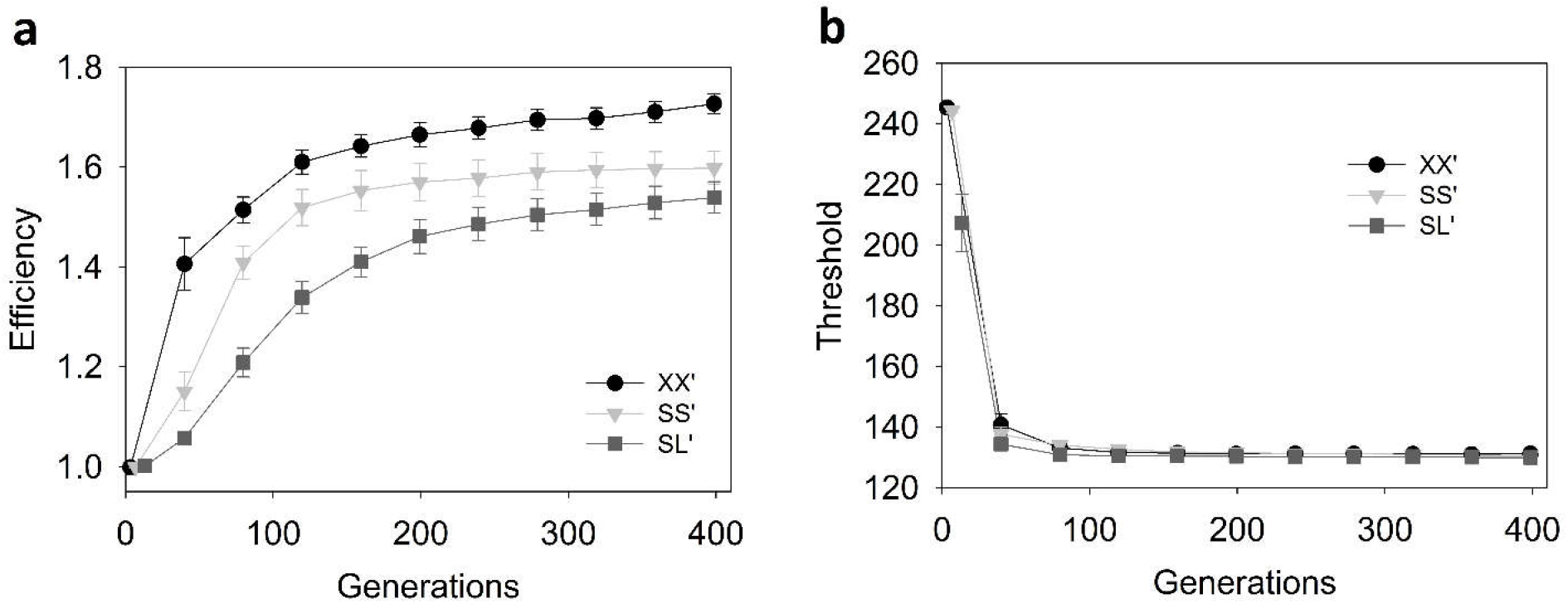
Adaptation in three populations with similar *HM* in terms of measures of fitness at the level of individuals. (**a**) Adaptive increase in averge individual efficiency within populations with similar harmonic mean size (mean ± SEM; 8 replicates). (**b**) Adaptive decrease in average individual threshold in populations with similar harmonic mean size (mean ± SEM; 8 replicates). Threshold evolved so quickly that its adaptive decrease did not refelct the differnce observed in *EoA* trjectories for *K* and *R* (Fig. 2 (Main text)). XX’: N_0_ ≈ 3.6*10^3^, bottleneck ratio: 1/10; SS’: N_0_ ≈ 1.8*10^3^, bottleneck ratio: 1/10^2^; SL’: N_0_ ≈ 9*10^2^, bottleneck ratio: 1/10^4^.

Multiple measures of fitness in our study revealed that harmonic mean is not a good predictor of adaptive trajectories because populations with similar harmonic mean size can have markedly different adaptive trajectories (Fig. S4 and Fig. 2 (Main-text)). Identical trends were observed when such populations (XX’, SS’, and SL’) were compared in terms of two different measures of population level fitness (Fig. 2 (Main-text)). In terms of fitness at the level of individuals, efficiency showed the same trend as *R* and *K* (Fig. S4a). However,the adaptive trajectories corresponding to XX’, SS’, and SL’ were almost identical when expressed in terms of threshold. Threshold evolved (decreased) so quickly and to such a large extent in almost all population types that we simulated in this study (regardless of their *HM*) that most populations had similar trajectories of threshold decrease (also see Fig. S7). Consequently, despite threshold being an important determinant of fitness, adaptive differences amongst populations were best expressed and explained in terms of trajectories of increase in efficiency and not in terms of decrease in threshold. The trends shown by adaptive trajectories of efficiency increase were identical to those shown by adaptive trajectories of *K* and *R*. Due to the above reasons, we focussed on population-wide trait distributions only in terms of efficiency.

##### Populations with similar HM show remarkable differences in distributions of efficiency

**Fig. S5.**
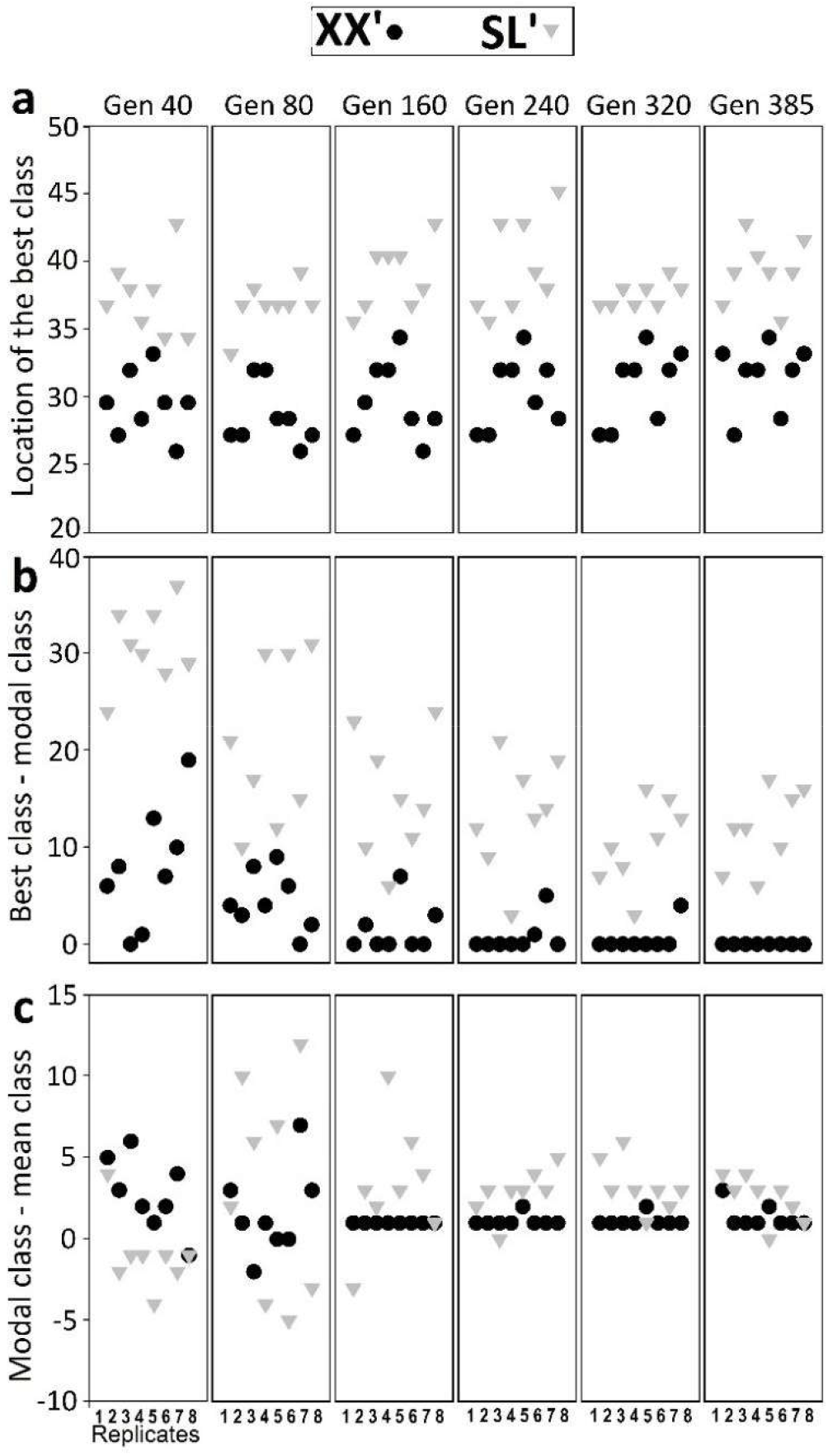
The distributions of efficiency across constituent individuals during adaptation in populations with similar HM. The individuals of each simulated population (8 replicate populations each of XX’ and SL’) were classified into to a discrete frequency distribution of their efficiency values (50 static classes) just prior to the bottleneck. Higher class indices correspond to higher efficiencies. The best phenotype (in terms of fitness) explored by SL’ was consistently fitter than the best phenotype explored by LB (**a**). The modal phenotype quickly converged with the best available phenotype in most XX’ populations but failed to do so in all SL’ populations (**b**). The mean phenotype in XX’ approached the best phenotype very closely (**b** and **c**). However, there was a consistently larger gap between the best phenotype and the modal phenotype in SL’ (**b**) and an even larger one between its best and mean phenotype (**b** and **c**). Our simulations revealed that populations with similar harmonic mean size can differ appreciably from each other not only in terms of their adaptive trajectories but also in terms of how the distribution of fitness amongst their constituent individuals changes during adaptation.

##### *EoA* shows an increasing relationship with *N_0_*

**Fig. S6.**
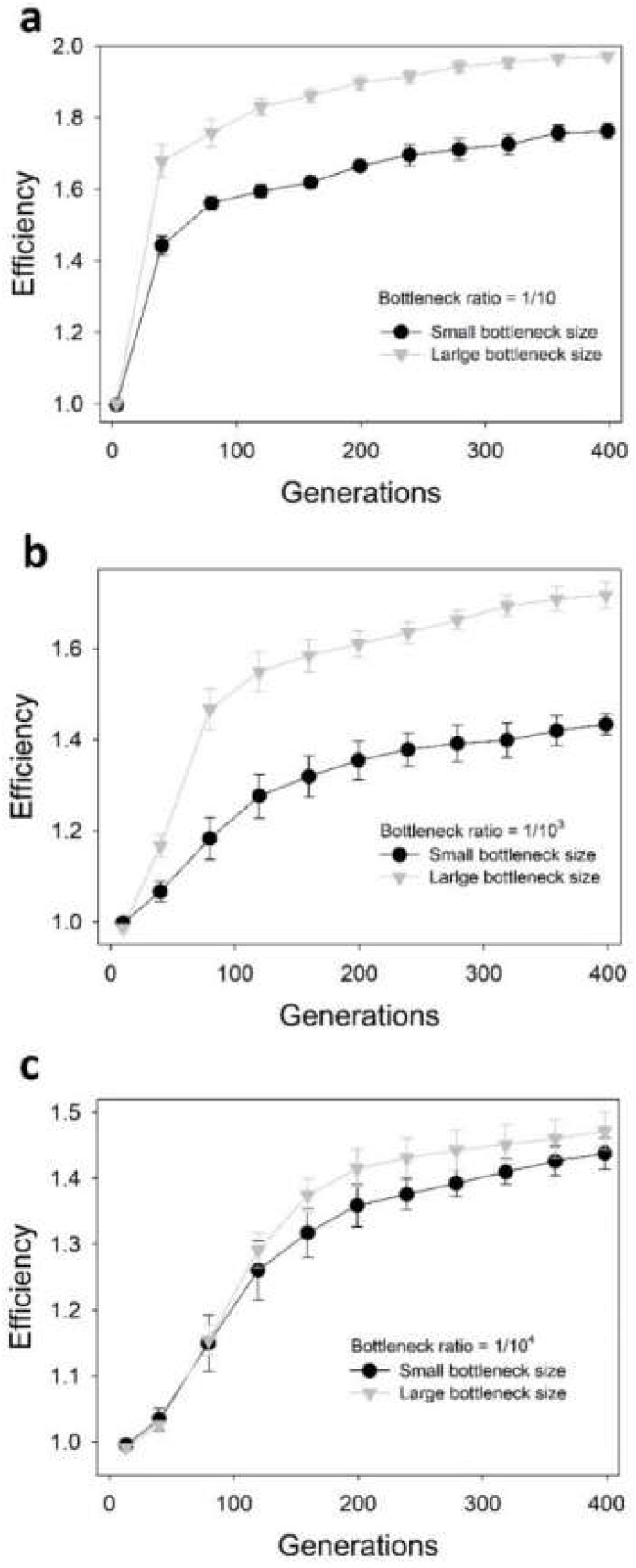
Simulations: *EoA* trajectories of populations with similar bottleneck ratio but different bottleneck sizes (*N_0_*) Data points show mean ± SEM; 8 replicates. (**a**) Bottleneck ratio = 1/10; Small bottleneck size (*N_0_*) ≈ 9000; Large bottleneck size (*N_0_*) ≈ 900000. (**b**) Bottleneck ratio = 1/10^3^; Small bottleneck size (*N_0_*) ≈ 90; Large bottleneck size (*N_0_*) ≈ 9000. (c) Bottleneck ratio = 1/10^4^; Small bottleneck size (*N_0_*) ≈ 90; Large bottleneck size (*N_0_*) ≈ 900.

##### *EoA* shows a decreasing relationship with *g*

**Fig. S7.**
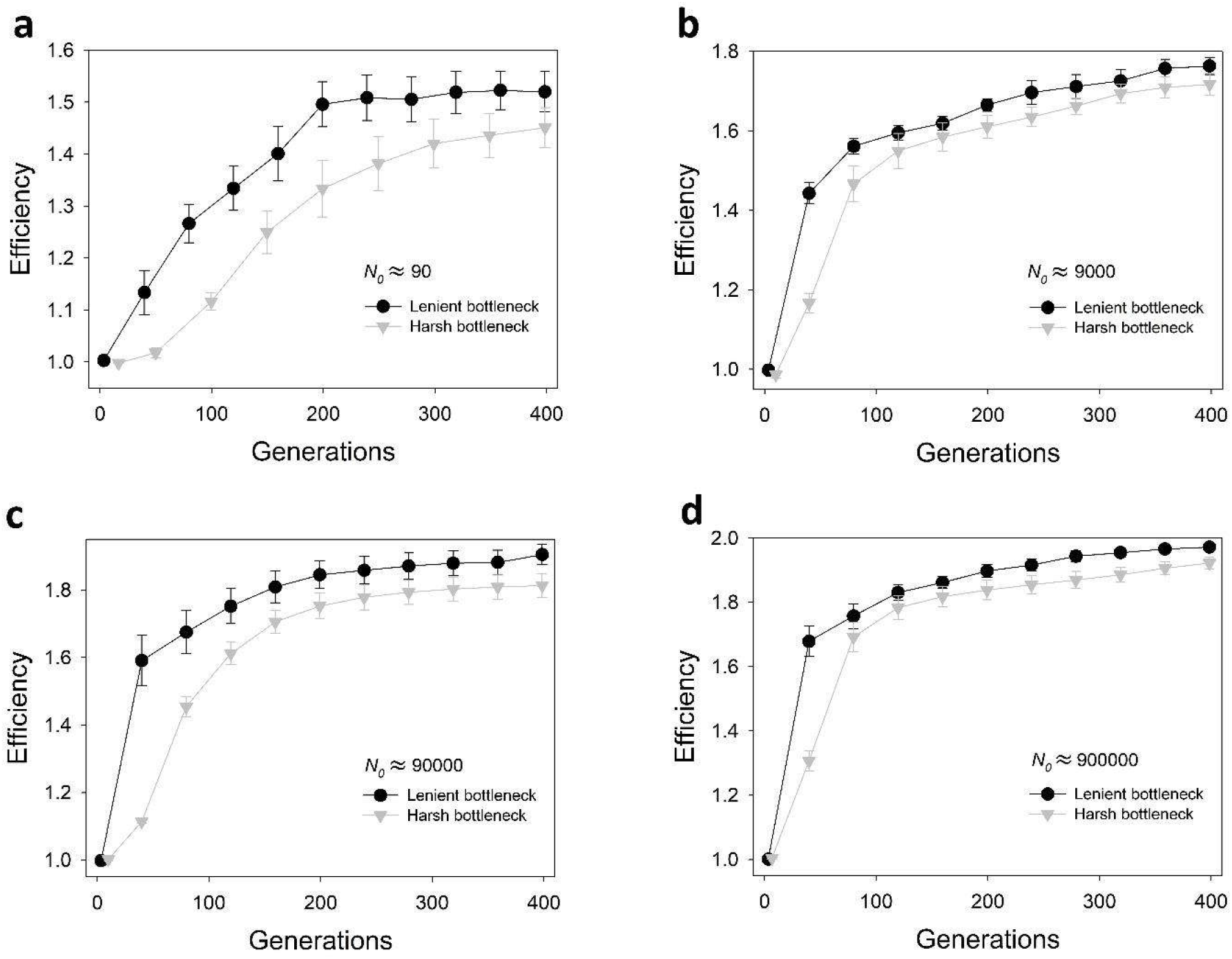
Simulations: *EoA* trajectories of populations with similar bottleneck size (*N_0_*) but different bottleneck ratios. Data points show mean ± SEM; 8 replicates. (**a**) *N_0_* ≈ 90; Lenient bottleneck = 1/10 (*g* = 3.32); Harsh bottleneck = 1/10^5^ (*g* = 16.61) (**b**) *N_0_* ≈ 9000; Lenient bottleneck = 1/10 (*g* = 3.32); Harsh bottleneck = 1/10^3^ (*g* = 9.96) (**c**) *N_0_* ≈ 90000; Lenient bottleneck = 1/10 (*g* = 3.32); Harsh bottleneck = 1/10^3^ (*g* = 9.96) (**d**) *N_0_* ≈ 900000; Lenient bottleneck = 1/10 (*g* = 3.32); Harsh bottleneck = 1/10^2^ (*g* = 6.64).

**Fig. S8.**
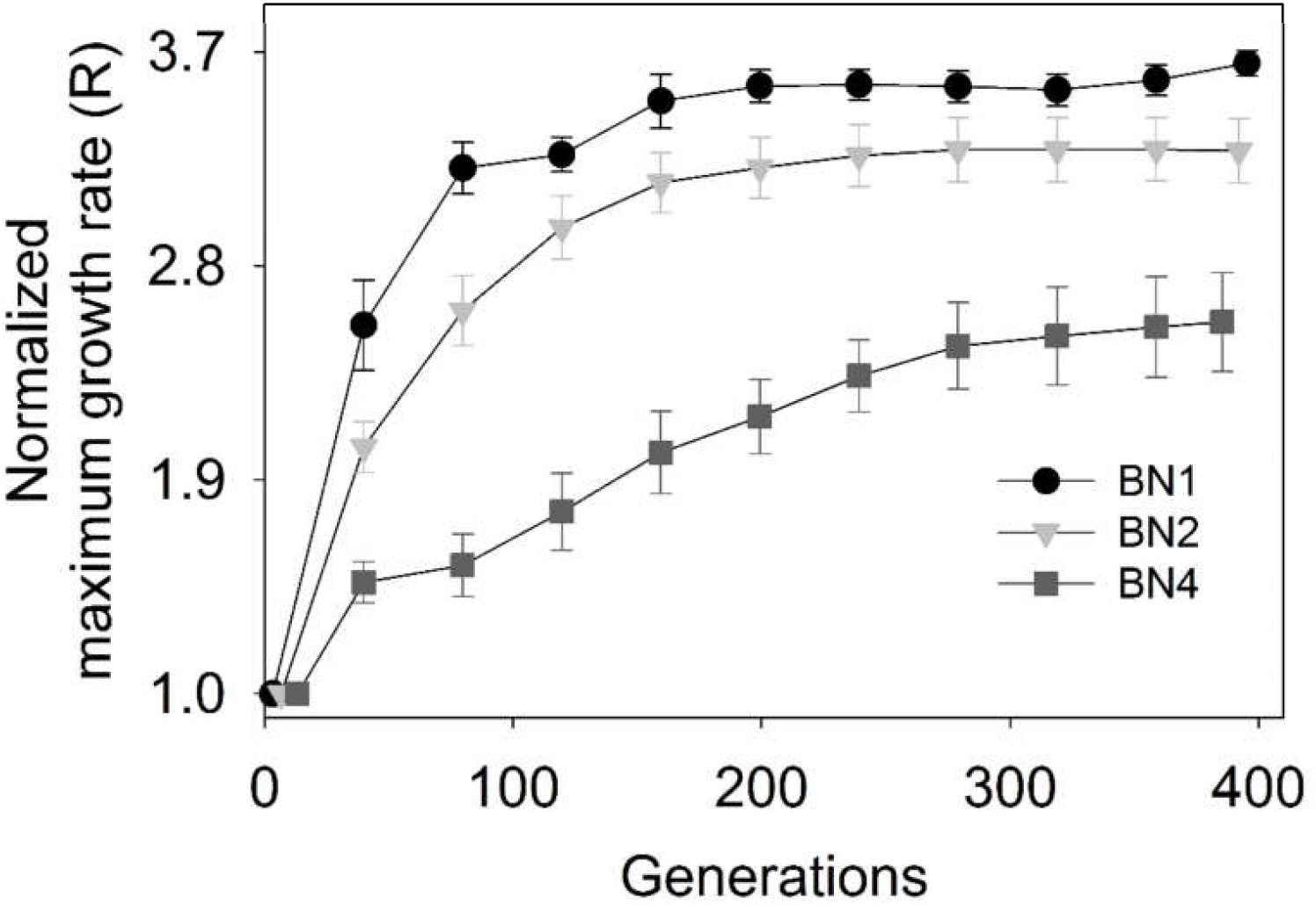
Simulations: *EoA* trajectories (in terms of *R*) of populations with similar bottleneck size (*N*_0_) but different bottleneck ratios. Data points show mean ± SEM; 8 replicates. All the population regimens shown here had *N_0_* ≈ 900. Bottleneck ratios: BN1: 1/10; BN2: 1/10^2^; BN4:1/10^4^. Larger values of *g* lead to reduced *EoA* for a given number of generations.

##### Threshold evolved (decreased) quickly and convergently in almost all populations

**Fig. S9.**
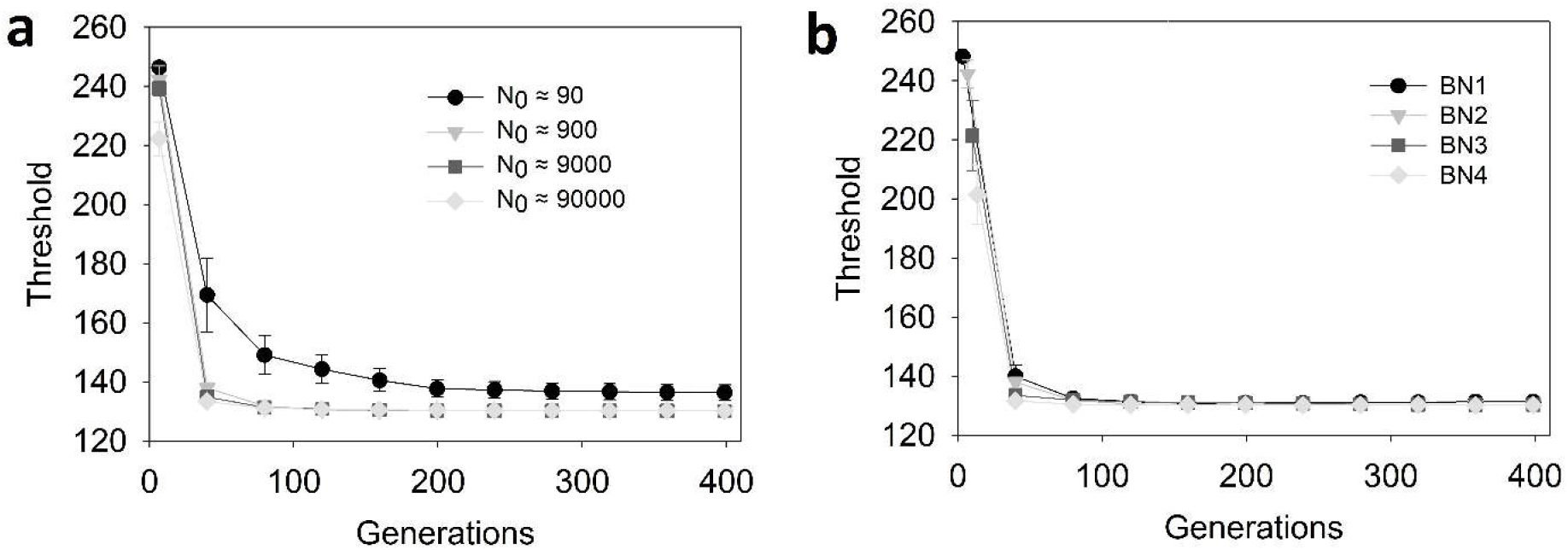
Rapid and convergent reduction in threshold. Threshold evolved (decreased) so quickly and convergently in most populations that we simulated that the effects The data points show mean ± SEM (N=8). The populations shown in (**a**) were bottlenecked 1/10^2^ periodically. The populations shown in (**b**) had N_0_ ≈ 900. Bottleneck ratios: BN1: 1/10; BN2: 1/10^2^; BN3: 1/10^3^; BN4:1/10^4^.

##### The relationship between *EoA* and *g* was robust to changes in mutation rates

**Fig. S10.**
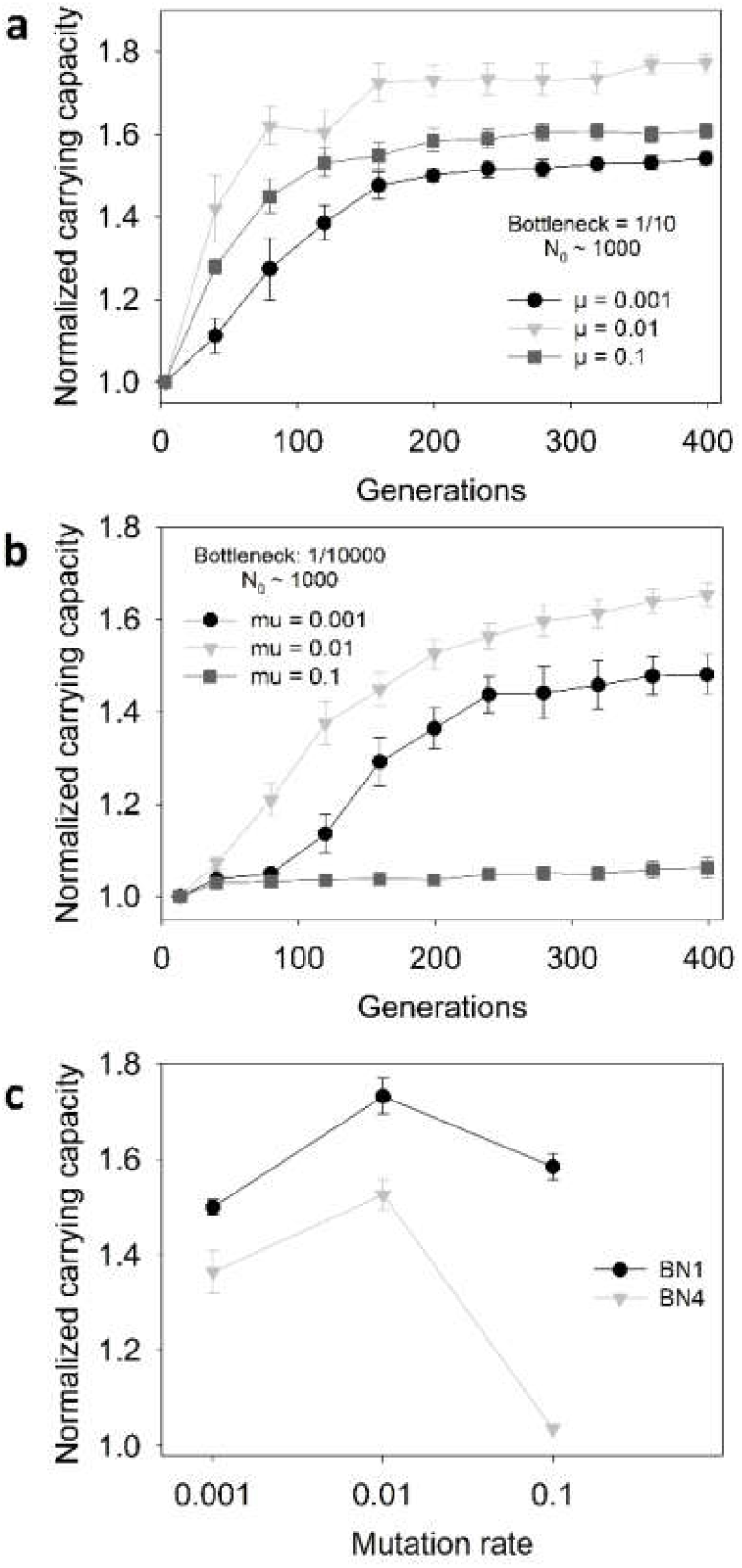
The relationship between *EoA* and *g* at three different mutation rates. (**a**) Adaptive increase in normalized carrying capacity in BN1 populations at three mutation rates. (**b**) Adaptive increase in normalized carrying capacity in BN4 populations at three mutation rates. (**c**) Normalized carrying capacity in BN1 and BN4 at generation 200 at three *µ* values. *EoA* exhibited a non-monotonous relationship with *µ* in both BN1 and BN4 populations, which is in line with theoretical expectations (Orr 2000). The negative dependence of *EoA* on *g* was robust to changes in mutation rate *(µ)* over a 100-fold range. We found that the relationship between *EoA* and *µ* can be influenced by bottleneck ratio. This is in agreement with recent empirical findings (Raynes et al. 2014; Raynes and Sniegowski 2014). The data points show mean ± SEM (8 replicates). Both BN1 and BN4 had similar bottleneck size *(N_0_* ≈ 900). BN1 experienced a periodic bottleneck of 1/10 whereas BN4 experienced a periodic bottleneck of 1/10^4^.

**Fig. S11.**
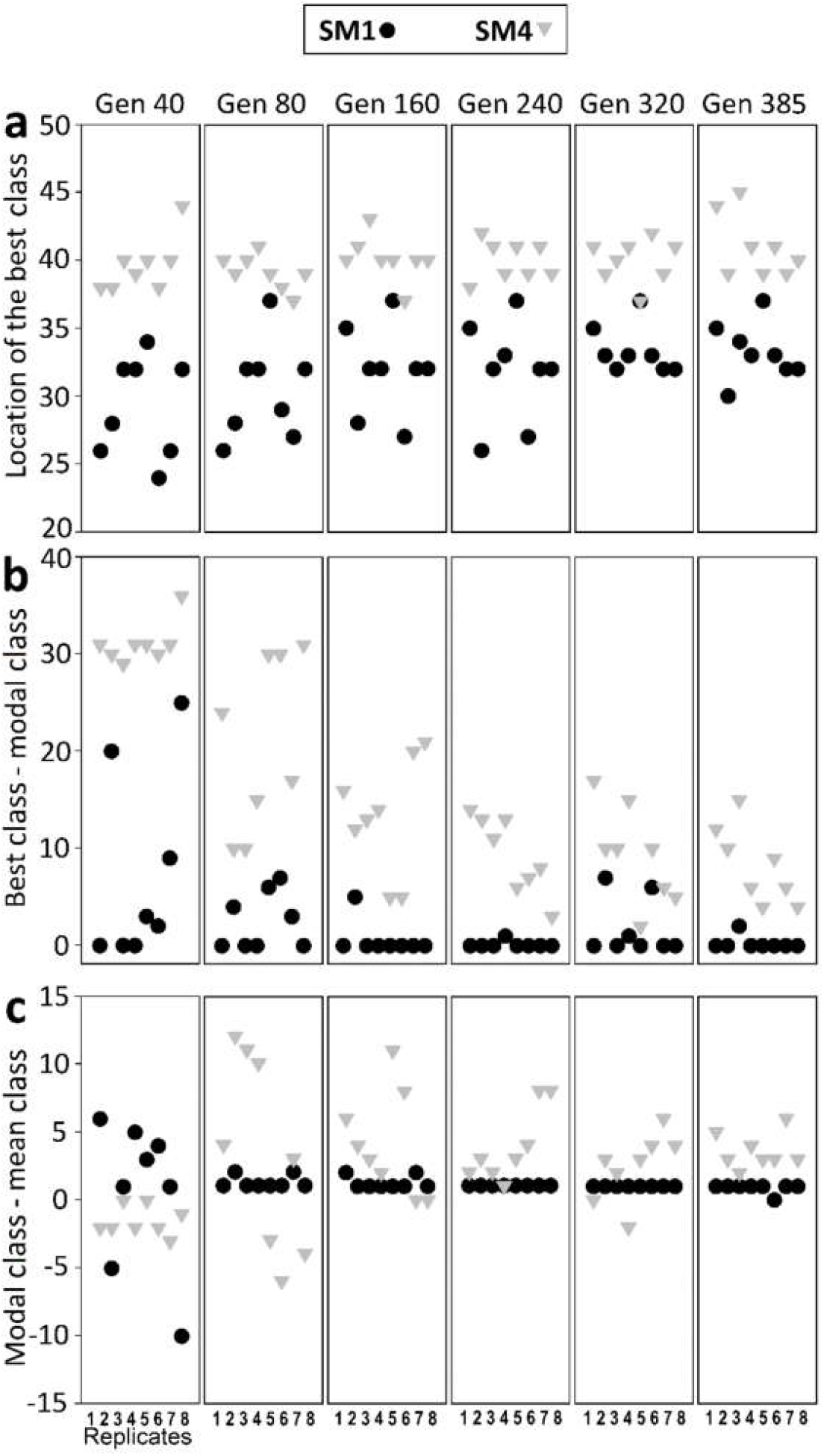
Simulations: Distributions of phenotypic effects across individuals during adaptation. The individuals of each simulated population (8 replicates each of SM1 and SM4) were classified into to a discrete frequency distribution of their efficiency values (50 static classes) prior to bottlenecks. Higher class indices correspond to higher efficiencies. (**a**) The best phenotype (in terms of efficiency) explored by SM4 was consistently fitter than the best phenotype explored by SM1. The modal phenotype quickly converged to the best available phenotype in all but one SM1 populations but failed to do so in all SM4 populations (**b**). The mean phenotype in SM1 approached the best phenotype very closely (**b and c**). However, there was a consistently larger gap between the best phenotype and the modal phenotype in SM4 (**c**) and an even larger one between its best and mean phenotype (**b and c**).

##### Locations of the best class before and after bottleneck

**Fig. S12.**
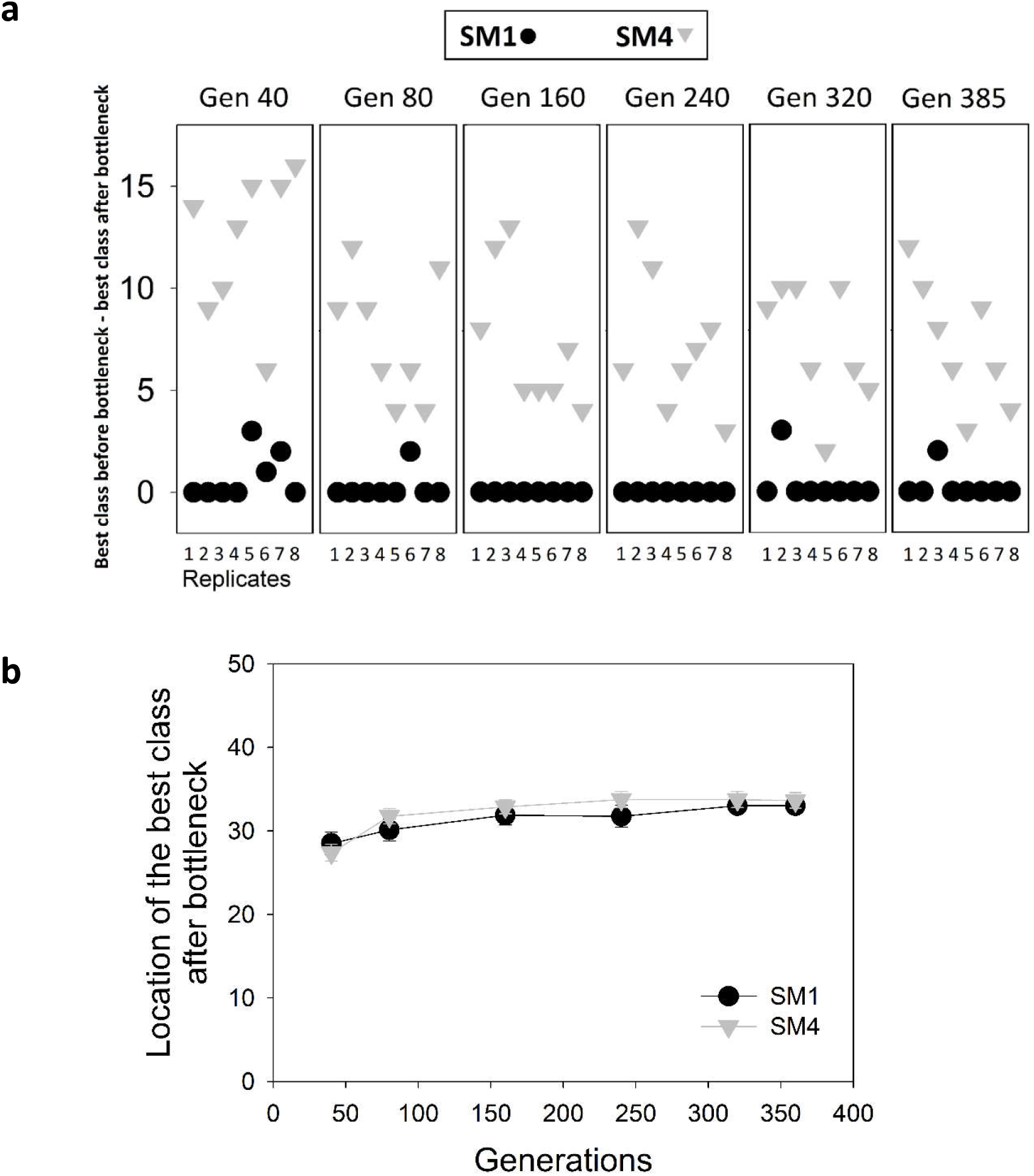
Locations of the best class before and after bottleneck. (**a**) Differences in the locations of the best class in the distribution of the efficeincy parameter before and after bottlenck in populations with similar *N_0_*. The individuals of each simulated population (8 replicate populations each of SM1 and SM4) were classified into to a discrete frequency distribution of their efficiency values (50 static classes). While the best class of SM1 could survive the bottleneck in most cases (black circles), the best class of SM4 invariably failed to survive its harsh bottleneck (grey triangles). (**b**) The locations of the best class after bottlenecks were remarkably similar across SM1 and SM4. The data points show mean ± SEM (8 replicates; some error bars are smaller than the data symbols.

Thus, SM4 populations will typically waste their best mutations (as the latter will be lost dring bottlenecks). However, SM1 populations will typically retain their best mutations across bottlenecks.

**Fig. S13.**
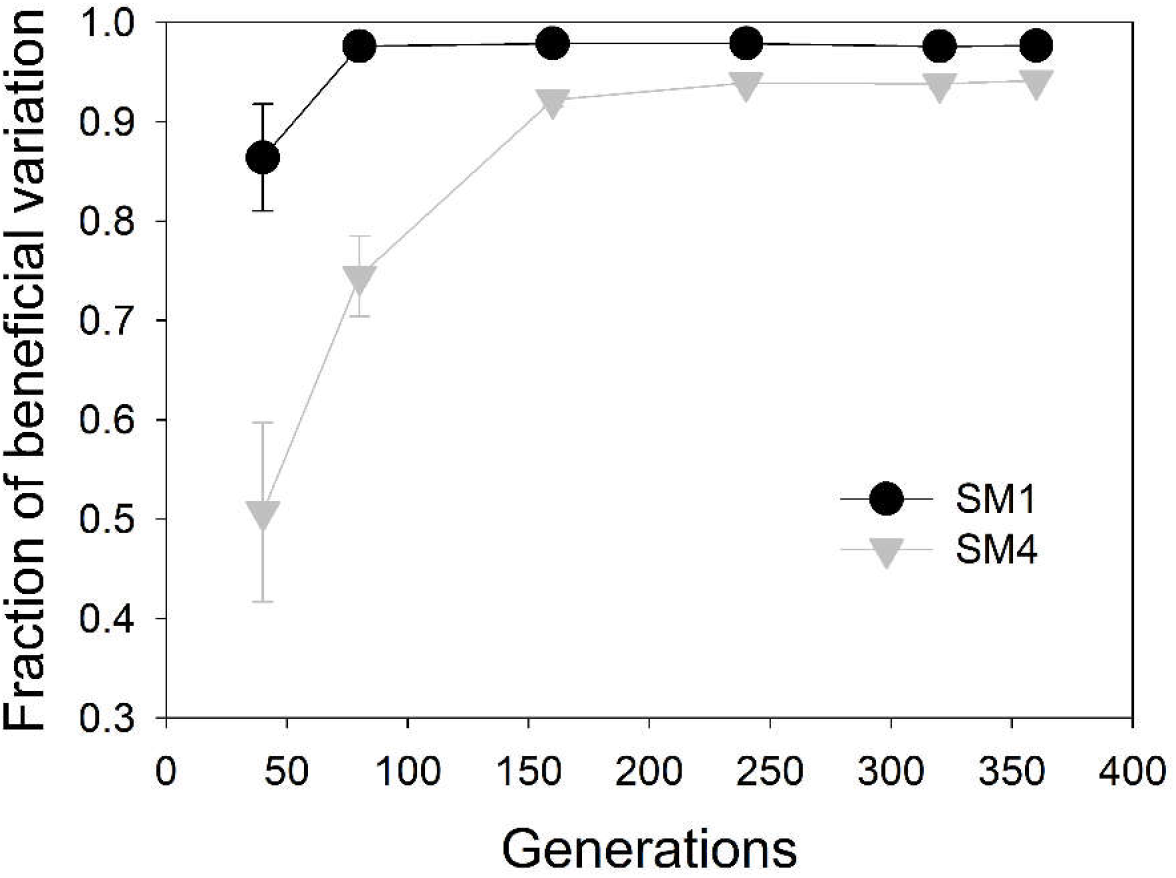
The fraction of beneficial variation with respect to the ancestor (in terms of efficiency) in SM1 and SM4. The data points show mean ± SEM (8 replicates; some error bars are smaller than the data symbols. Beneficial variation is refers to the variation by the individuals that are fitter than the wild-type (ancestor).

##### *N_0_/g* can predict *EoA* better than *N_0_g*

**Fig. S14.**
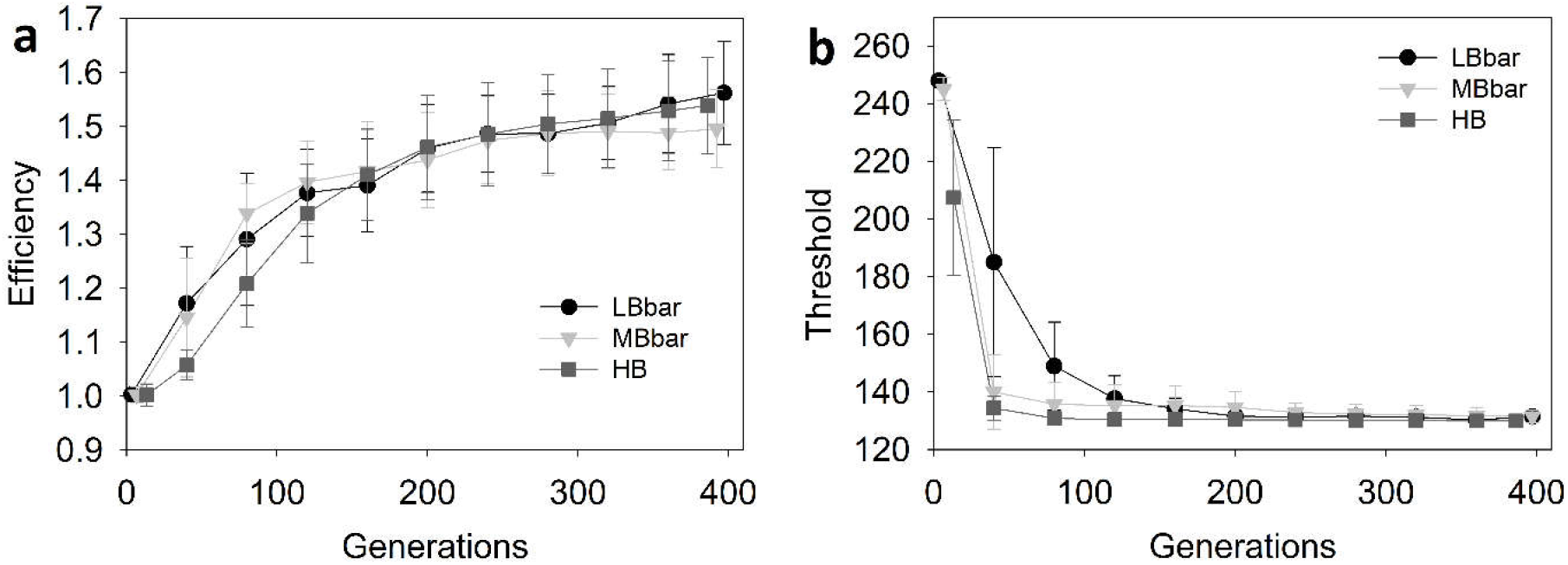
*N_0_/g* was a better predictor of *EoA* than *N_0_g*. Adaptive trajectories in populations with similar *N_0_/g* expressed in terms of individual-level fitness parameters: efficiency (**a**), threshold (**b**). The data points show mean ± SD (8 replicates) in (**a**) and (**b**). Populations with similar *N_0_/g* (LBbar and HB) match more closely in terms of mean adaptive trajectories than populations with similar *N_0_g* (LB and HB). LBbar: *N_0_* ≈ 225; bottleneck ratio = 1/10; MBbar: *N_0_* ≈ 450; bottleneck ratio = 1/10^2^; *N_0_* ≈ 900; bottleneck ratio = 1/10^4^. HB: *N_0_* ≈ 900, bottleneck ratio: 1/10^4^;

Populations with similar *N_0_/g* had remarkably similar adaptive trajectories in terms of both efficiency and threshold. These populations had similar adaptive trajectories despite differing in terms of the intensity of the periodic botleneck over a 1000-fold range. While *N_0_*g* failed to predict adaptive trajectories over this bottleneck range (Fig. 2 (Main text)), *N_0_/g* could act as a much better predictor of adaptive trajectories.

**Fig. S15.**
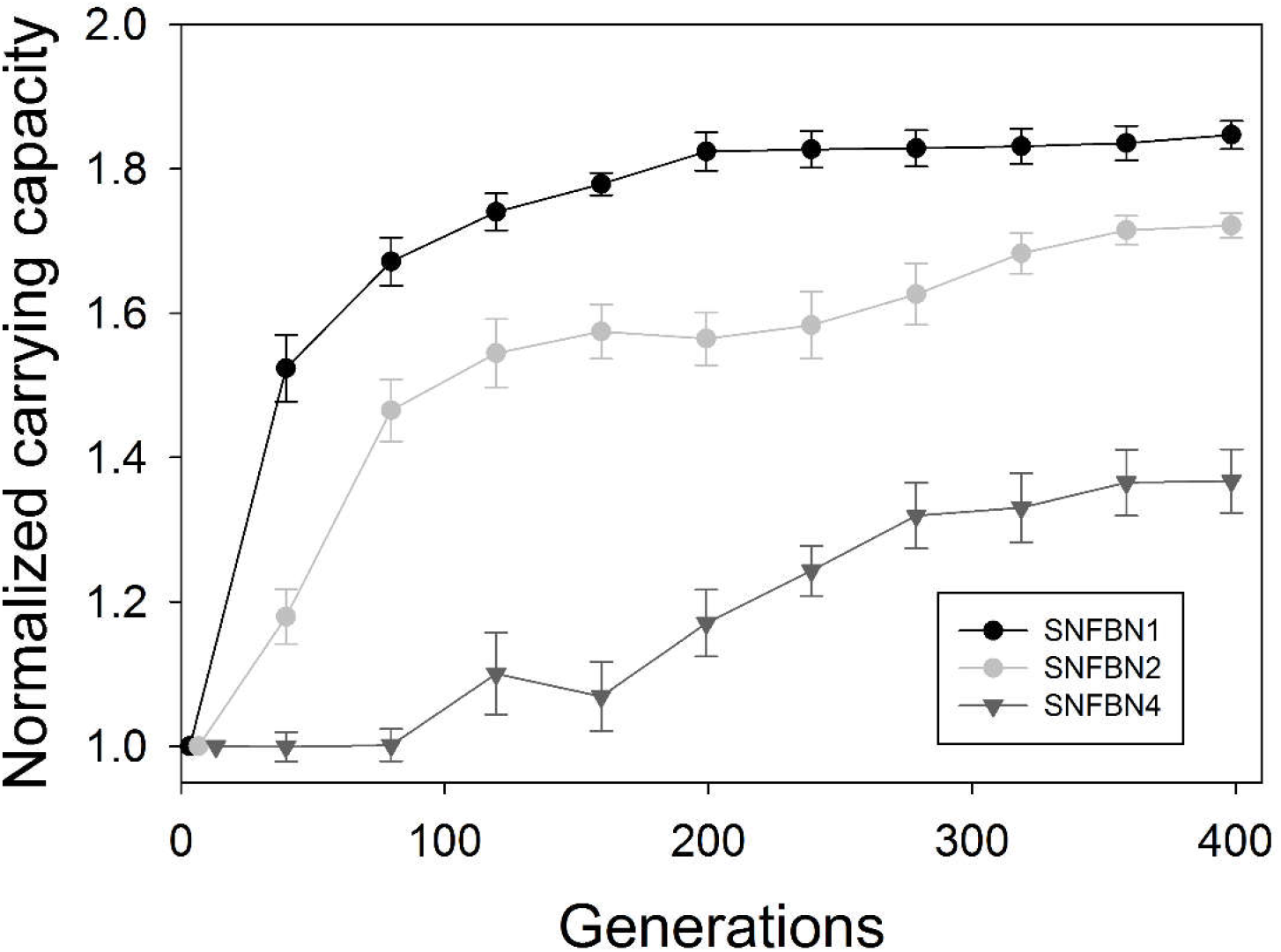
*EoA* in populations with similar initial *N_f_*, but different *N_0_g*. When initial *N_f_* is constant, populations with higher *N_0_g* also have higher *N_0_/g*, and thus show higher *EoA*. However, when *N_f_* is not constant (e.g., see Figs. 2, 3b, 4, and 5 of the main text), higher *N_0_g* does not lead to higher *EoA*. SNFBN1: *N_0_* ≈ 9000; bottleneck ratio = 1/10; *N_0_g* ≈ 29900; *N_0_/g* ≈ 2711. SNFBN2: *N_0_* ≈ 900; bottleneck ratio = 1/10^2^; *N_0_g* ≈ 5975; *N_0_/g* ≈ 136. SNFBN4: *N_0_* ≈ 9; bottleneck ratio = 1/10^4^; *N_0_g* ≈ 120; *N_0_/g* ≈ 0.68.

The above graph shows that both *N_0_/g* and *N_0_g* can successfully predict *EoA* if the populations under consideration have similar initial *N_f_*.

## References

Abdi, H. (2010). Holm’s sequential Bonferroni procedure. Encyclopedia of research design, 1–8.

Abel, S., Abel zur Wiesch, P., Davis, B. M., & Waldor, M. K. (2015). Analysis of Bottlenecks in Experimental Models of Infection. PLoS Pathog, 11(6), e1004823. doi:10.1371/journal.ppat.1004823

Barrick, J. E., Yu, D. S., Yoon, S. H., Jeong, H., Oh, T. K., Schneider, D., et al. (2009). Genome evolution and adaptation in a long-term experiment with Escherichia coli. Nature, 461(7268), 1243–1247. doi:10.1038/nature08480

Campos, P. R. A., & Wahl, L. M. (2009). The Effects of Population Bottlenecks on Clonal Interference, and the Adaptation Effective Population Size. Evolution, 63(4), 950–958. doi:10.1111/j.1558-5646.2008.00595.x

Campos, P. R. A., & Wahl, L. M. (2010). The Adaptation Rate of Asexuals: Deleterious Mutations, Clonal Interference and Population Bottlenecks. Evolution, 64(7), 1973–1983. doi:10.1111/j.1558-5646.2010.00981.x

Charlesworth, B. (2009). Effective population size and patterns of molecular evolution and variation. Nature Reviews Genetics, 10(3), 195–205.

Cohen, J. (1988). Statistical power analysis for the behavioral sciences. Hillsdale, N.J.: L. Erlbaum Associates.

Crow, J. F. (1958). Some possibilities for measuring selection intensities in man. Human Biology, 30(1), 1–13.

De Visser, J. a. G. M., & Rozen, D. E. (2005). Limits to adaptation in asexual populations. Journal of Evolutionary Biology, 18(4), 779–788. doi:10.1111/j.1420-9101.2005.00879.x

Desai, M. M., & Fisher, D. S. (2007). Beneficial mutation–selection balance and the effect of linkage on positive selection. Genetics, 176(3), 1759–1798.

Desai, M. M., Fisher, D. S., & Murray, A. W. (2007). The Speed of Evolution and Maintenance of Variation in Asexual Populations. Current Biology, 17(5), 385–394. doi:10.1016/j.cub.2007.01.072

Gerrish, P. J., & Lenski, R. E. (1998). The fate of competing beneficial mutations in an asexual population. Genetica, 102–103(0), 127–144. doi:10.1023/A:1017067816551

Heffernan, J. M., & Wahl, L. M. (2002). The effects of genetic drift in experimental evolution. Theoretical Population Biology, 62(4), 349–356. doi:10.1016/S0040-5809(02)00002-3

Karve, S. M., Bhave, D., Nevgi, D., & Dey, S. (2016). Escherichia coli populations adapt to complex, unpredictable fluctuations by minimizing trade-offs across environments. Journal of Evolutionary Biology, 29(12), 2545–2555. doi:10.1111/jeb.12972

Karve, S. M., Daniel, S., Chavhan, Y. D., Anand, A., Kharola, S. S., & Dey, S. (2015). Escherichia coli populations in unpredictably fluctuating environments evolve to face novel stresses through enhanced efflux activity. Journal of Evolutionary Biology, 28(5), 1131–1143. doi:10.1111/jeb.12640

Kassen, R. (2014). Experimental evolution and the nature of biodiversity.

Katju, V., Packard, L. B., Bu, L., Keightley, P. D., & Bergthorsson, U. (2015). Fitness decline in spontaneous mutation accumulation lines of Caenorhabditis elegans with varying effective population sizes. Evolution; International Journal of Organic Evolution, 69(1), 104–116. doi:10.1111/evo.12554

Kawecki, T. J., Lenski, R. E., Ebert, D., Hollis, B., Olivieri, I., & Whitlock, M. C. (2012). Experimental evolution. Trends in Ecology & Evolution, 27(10), 547–560. doi:10.1016/j.tree.2012.06.001

Kimura, M. (1983). The neutral theory of molecular evolution. Cambridge University Press. https://books.google.co.in/books?hl=en&lr=&id=olIosumpevyc&oi=fnd&pg=PR9&dq=the+neutral+theory+of+molecular+evolution&ots=P-u4qegocd&sig=Pnbi0shzvvrachlhrrdkn_i38a. Accessed 8 February 2017

LaBar, T., & Adami, C. (2016). Different Evolutionary Paths to Complexity for Small and Large Populations of Digital Organisms. PLOS Computational Biology, 12(12), e1005066. doi:10.1371/journal.pcbi.1005066

Lachapelle, J., Reid, J., & Colegrave, N. (2015). Repeatability of adaptation in experimental populations of different sizes. Proceedings of the Royal Society of London B: Biological Sciences, 282(1805), 20143033. doi:10.1098/rspb.2014.3033

Lanfear, R., Kokko, H., & Eyre-Walker, A. (2014). Population size and the rate of evolution. Trends in Ecology & Evolution, 29(1), 33–41. doi:10.1016/j.tree.2013.09.009

Lenski, R. E., Rose, M. R., Simpson, S. C., & Tadler, S. C. (1991). Long-Term Experimental Evolution in Escherichia coli. I. Adaptation and Divergence During 2,000 Generations. The American Naturalist, 138(6), 1315–1341.

Novak, M., Pfeiffer, T., Lenski, R. E., Sauer, U., & Bonhoeffer, S. (2006). Experimental Tests for an Evolutionary Trade‐Off between Growth Rate and Yield in E. coli. The American Naturalist, 168(2), 242–251. doi:10.1086/an.2006.168.issue-2

Patwa, Z., & Wahl, L.. (2008). The fixation probability of beneficial mutations. Journal of the Royal Society Interface, 5(28), 1279–1289. doi:10.1098/rsif.2008.0248

Petit, N., & Barbadilla, A. (2009). Selection efficiency and effective population size in Drosophila species. Journal of Evolutionary Biology, 22(3), 515–526. doi:10.1111/j.1420-9101.2008.01672.x

Ramsayer, J., Kaltz, O., & Hochberg, M. E. (2013). Evolutionary rescue in populations of Pseudomonas fluorescens across an antibiotic gradient. Evolutionary Applications, 6(4), 608–616. doi:10.1111/eva.12046

Raynes, Y., Halstead, A. L., & Sniegowski, P. D. (2014). The effect of population bottlenecks on mutation rate evolution in asexual populations. Journal of Evolutionary Biology, 27(1), 161–169. doi:10.1111/jeb.12284

Raynes, Yevgeniy, Gazzara, M. R., & Sniegowski, P. D. (2012). Contrasting Dynamics of a Mutator Allele in Asexual Populations of Differing Size. Evolution, 66(7), 2329–2334. doi:10.1111/j.1558-5646.2011.01577.x

Rice, S. H. (2004). Evolutionary theory: mathematical and conceptual foundations. Sinauer Associates.

Rozen, D. E., de Visser, J. A. G. M., & Gerrish, P. J. (2002). Fitness Effects of Fixed Beneficial Mutations in Microbial Populations. Current Biology, 12(12), 1040–1045. doi:10.1016/S0960-9822(02)00896-5

Rozen, D. E., Habets, M. G. J. L., Handel, A., & de Visser, J. A. G. M. (2008). Heterogeneous Adaptive Trajectories of Small Populations on Complex Fitness Landscapes. PLoS ONE, 3(3), e1715. doi:10.1371/journal.pone.0001715

Samani, P., & Bell, G. (2010). Adaptation of experimental yeast populations to stressful conditions in relation to population size. Journal of Evolutionary Biology, 23(4), 791–796. doi:10.1111/j.1420-9101.2010.01945.x

Sniegowski, P. D., & Gerrish, P. J. (2010). Beneficial mutations and the dynamics of adaptation in asexual populations. Philosophical Transactions of the Royal Society B: Biological Sciences, 365(1544), 1255–1263. doi:10.1098/rstb.2009.0290

Szendro, I. G., Franke, J., Visser, J. A. G. M. de, & Krug, J. (2013). Predictability of evolution depends nonmonotonically on population size. Proceedings of the National Academy of Sciences, 110(2), 571–576. doi:10.1073/pnas.1213613110

Vogwill, T., Phillips, R. L., & Gifford, D. R. (2016). Divergent evolution peaks under intermediate population bottlenecks during bacterial experimental evolution. Proc. R. Soc. B, 283(1835), 20160749. doi:10.1098/rspb.2016.0749

Wahl, L. M., & Gerrish, P. J. (2001). The Probability That Beneficial Mutations Are Lost in Populations with Periodic Bottlenecks. Evolution, 55(12), 2606–2610. doi:10.1111/j.0014-3820.2001.tb00772.x

Wahl, L. M., Gerrish, P. J., & Saika-Voivod, I. (2002). Evaluating the impact of population bottlenecks in experimental evolution. Genetics, 162(2), 961–971.

Wahl, L. M., & Zhu, A. D. (2015). Survival probability of beneficial mutations in bacterial batch culture. Genetics, 200(1), 309–320. doi:10.1534/genetics.114.172890

Wang, J., Santiago, E., & Caballero, A. (2016). Prediction and estimation of effective population size. Heredity, 117(4), 193–206. doi:10.1038/hdy.2016.43

Wilke, C. O. (2004). The Speed of Adaptation in Large Asexual Populations. Genetics, 167(4), 2045–2053. doi:10.1534/genetics.104.027136

## References to Supplementary Information

Abdi, H. (2010). Holm’s sequential Bonferroni procedure. Encyclopedia of research design, 1–8.

Elena, S. F., Cooper, V. S., & Lenski, R. E. (1996). Punctuated Evolution Caused by Selection of Rare Beneficial Mutations. Science, 272(5269), 1802–1804. doi:10.1126/science.272.5269.1802

Orr, H. A. (2000). The Rate of Adaptation in Asexuals. Genetics, 155(2), 961–968.

Raynes, Y., & Sniegowski, P. D. (2014). Experimental evolution and the dynamics of genomic mutation rate modifiers. Heredity, 113(5), 375–380. doi:10.1038/hdy.2014.49

Tenaillon, O., Barrick, J. E., Ribeck, N., Deatherage, D. E., Blanchard, J. L., Dasgupta, A., et al. (2016). Tempo and mode of genome evolution in a 50,000-generation experiment. Nature, 535. http://www.biorxiv.org/content/early/2016/01/15/036806.abstract. Accessed 3 November 2016

